# Plant PhysioSpace: a robust tool to compare stress response across plant species

**DOI:** 10.1101/2020.11.16.384305

**Authors:** Ali Hadizadeh Esfahani, Janina Maß, Asis Hallab, Bernhard M. Schuldt, David Nevarez, Björn Usadel, Mark-Christoph Ott, Benjamin Buer, Andreas Schuppert

## Abstract

Generalization of transcriptomics results can be achieved by comparison across experiments, which is based on integration of interrelated transcriptomics studies into a compendium. Both characterization of the fate of the organism as well as distinguishing between generic and specific responses can be gained in such a broader context. There are numerous methods for analyzing such data sets, most focusing on gene-wise dimension reduction to obtain marker genes and gene sets, e.g. for pathway analysis. Relying only on isolated biological modules might lead to missing of important confounders and relevant context.

We have developed a novel method called Plant PhysioSpace, which provides the ability to compute experimental conditions across species and platforms without a priori reducing the reference information to specific gene-sets. It extracts physiologically relevant signatures from a reference data set, a collection of public data sets, by integrating and transforming heterogeneous reference gene expression data into a set of physiology-specific patterns. New experimental data can be mapped to these patterns, resulting in similarity scores which provide quantitative likeness of the new experiment to the a priori compendium.

Because of its robustness against noise and platform bias, Plant PhysioSpace can be used as an inter-species or cross-platform similarity measure. We have demonstrated its success in translating stress responses between different species and platforms (including single-cell technologies).

We also report the implementation of two R packages, one software and one data package, and a shiny web application, which provides plant biologists convenient ways to access our method and precomputed models from four different species.

## 1 Introduction

As a consequence of their non-motile nature, plants developed a peculiarly organized yet labyrinthine response system to external biotic and abiotic stresses. Exploiting this complex system has been playing an important role in achieving sustainable plant protection in agriculture. Instances of tweaking the plant defense system for obtaining better crops are numerous. For instance, priming, i.e. promoting plants to a primed state of defense, has been known, investigated and utilized for decades if not centuries [1, 2]. By exposure to biotic stresses (e.g. microbe-, pathogen-, herbivore-associated molecular patterns) or abiotic stresses (for instance harsh temperatures, drought or damage-associated molecular patterns), plants switch to a primed reinforced defense state. In this primed state, they can display sharper stress response, which in turn results in more robust and resilient organisms. By artificially exposing plants to biotic and abiotic stresses directly, or to some natural or synthetic chemicals which provoke the same defense response, it is possible to engineer tougher plants [3]. Another example of crop engineering is by genetically modifying (GM) plants to attain higher tolerance to stress [4]. Introducing a single gene encoding C-5 sterol desaturase (FvC5SD) from *Collybia velutipes* to tomato is an instance of GM crop research, and it brings about a drought-tolerant and fungal resistant crop [5, 6]. Obtaining resistance to papaya ringspot virus (PRSV) in transgenic papaya is another famous example. The resistant papaya gains the protection by expressing the PSRV coat protein transgene [7].

In research experiments aiming to modify the plant’s defense system, such as the examples mentioned above, the stress responses of plants under study are to be thoroughly examined and contrasted to wild types. We argue that a tool, which is capable of quantitatively and dependably measuring the speed and intensity of stress responses in plants, can be of great assistance in this field of research. Hence, we present Plant PhysioSpace, an advanced computational tool based on PhysioSpace [8], for quantitative analysis of stress responses in plants.

Sequencing technologies are commonly used for studying the changes in the plants under examination. However, analysis of the results mostly focuses on gene-wise dimension reduction of data to obtain a list of genes, with the rest of the analysis pipeline fixating on the genes in the list. By design, Plant PhysioSpace extracts physiologically relevant information out of intricately convoluted gene expression data without reducing dimensions, providing a direct link from sequencing data to physiological processes. Since it is computationally cheap, the tool is able to train on a vast amount of retrospectively available data, allowing explicit integration of established knowledge and data, eventuating in robust results when testing the method on small data sets generated in specific experiments.

Plant PhysioSpace comprises two compartments: the space generation, the algorithm which elicits information from big data, and the physio-mapping, the process with which new data can be analyzed by comparison to the extracted information (Fig. 1). Compared to the machine learning nomenclature, space generation is analogous to *training* and physio-mapping to *testing*.

**Figure 1:**
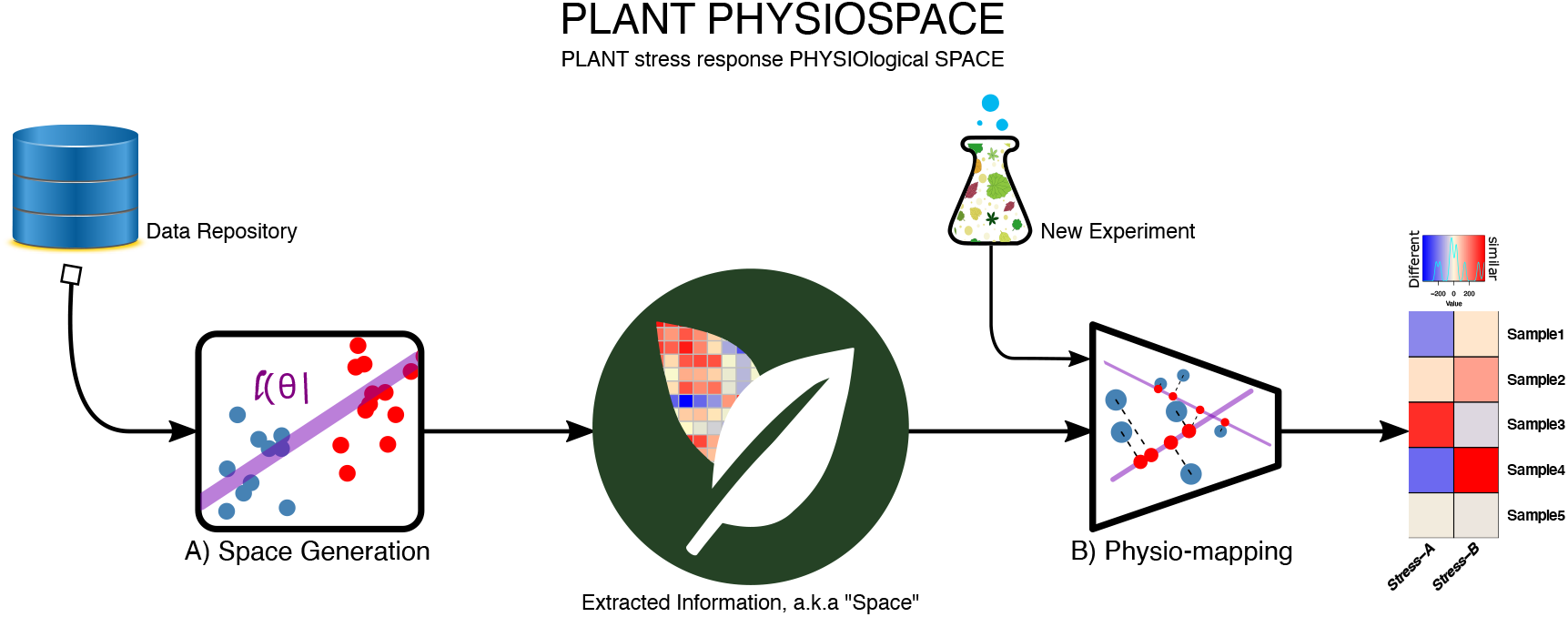
Plant PhysioSpace Overview. The method consists of two main sections: space generation and Physiomapping. In space generation (A), data from public repositories is processed and its information is extracted. After trimming, the extracted information is stored in matrices called “space”, representing physiology-relevant expression patterns. Physio-mapping (B) uses a space to analyze new, unknown data, for example from a new experiment. The new data is mapped to the generated space, resulting in “similarity” scores that indicate the likeness of the new data to the known physiological processes.

In this study, we focused on the application of our novel method in stress response analysis. As one of the fiercest adversaries of plants, biotic and abiotic stresses take a toll on commercial agriculture. Plant PhysioSpace can aid in engineering impervious crops, by quantitatively analyzing the effect of a new mutation or treatment on plant’s resistance.

Another long-lasting question in the field of stress response research is the potential heterogeneity in response among cell types under the stress. Generally, sequencing is done on thousands to millions of cells, revealing only the average effect on the bulk tissue, thus lacking the direct assessment of cells. The individual and unique role of cell types shape the function of their parent tissue. Hence, by careful examination of stressed tissue cells, the difference between their behavior, and the in-between interplay among them, one can gain new insights into the complex mechanisms shaping the plant stress response.

Since 2009, more and more single cell data sets are becoming available publicly. As with other new technologies, the focus is mainly on human and animal tissue sequencing. Lack of data availability is especially true for plant studies on account of processing tissues with cell walls has been a bothersome obstacle for single cell technologies, as they mostly result in low capture rates. But recent leaps in single cell sequencing technologies, e.g. the 10X platform, increased the resolution of single cell data, eventuating in a few plant single cell experiments [9, 10, 11, 12, 13]. Mostly, scRNA-seq studies follow the same analysis pipeline [14, 15]. In a nutshell, the highest variable genes are selected and gone through principal component analysis (PCA) and t-distributed stochastic neighborhood embedding (tSNE) or uniform manifold approximation and projection (UMAP) to demonstrate the underlying structure. Subsequently, Clustering or regression algorithms are used to identify biologically relevant groups (e.g. groups of cells with a similar response), or trends (e.g. pseudotemporal axis of cell development), from the underlying data structure [16]. Although such technologies as 10X made plant single cell sequencing possible, they are far from perfect. For instance, compared to bulk sequencing technologies, technical noise has a higher interference on single cell reads, which calls for developing sophisticated bioinformatic analysis tools to handle those interferences.

This paper has been organized in the following way: It begins with a brief explanation of the Plant PhysioSpace algorithm, which includes a review of the already published method, plus the modifications adapting the method to the field of plant stress research. The paper will then go on to the benchmarking section, in which the method’s performance is assessed in translating stress response among different experiments, platforms, and species. Benchmarking is followed by two application showcases, in which we demonstrate two Plant PhysioSpace use-cases: investigating time-series data from biotic-stressed wheat, and analyzing a heat-stressed single cell data set. Finally, the discussion gives a brief summary and critique of the findings.

## 2 Materials and Methods

### 2.1 Data Preparation

While setting up a PhysioSpace matrix, our method requires extensive training data for achieving adequate robustness. This training data can be retrieved from retrospective data sets. To that end, we curated more than 4000 plant stress response gene expression samples from GEO^1^ and SRA^2^. More specifically, 2480 *A. thaliana (Arabidopsis thaliana*) array samples, 967 *A. thaliana* RNA-seq samples, 146 *Oryza sativa* array samples, 172 *Glycine max* array samples, and 104 *Triticum aestivum* array samples were used for space generation. Each sample is annotated with a label from a stress set. In this study, samples are divided into Aluminum, Magnesium, Biotic, Cold, Drought, FarRed, FeDeficiency, Genotoxic, Heat, Herbicide, Hormone, Hypoxia, Light, LowPH, Metabolic, Mutant, Nitrogen, Osmotic, Radiation, Salt, Submergence, UV and Wounding stress groups. For samples which underwent more than one stress, new labels were generated by concatenating existing labels from the stress set. For example, ‘Biotic.Drought’ designates a sample which sustained both Biotic and Drought stresses.

Samples corresponding to each species are normalized in bulk to remove the batch effect. We used robust multi-array average or RMA [17] for normalizing microarray RAW data files and a pipeline consisting of Fastq-dump, Trimmomatic [18], Star aligner [19] and featureCounts [20] to derive counts from SRA records.

### 2.2 PhysioSpace Method

**PhysioSpace** is a supervised dimension reduction method, which aims to extract relevant **physio**logical information from big data sets and store it in a mathematical **space** [8]. The method can be divided into two main steps: space generation and physio-mapping (Fig. 1).

#### 2.2.1 Space Generation

After preparing the data, we derive the physiologically relevant information from normalized data and store this information in a mathematical space. This step is comparable to *training* in machine learning terminology. Space generation is done in two stages: space extraction and space trimming. The former stage is identical to the method described previously in [8]. However, the latter, space trimming, is a novel addition for adapting the method for studying plant stress response.

##### Space Extraction

In the PhysioSpace method, all samples are analyzed contrastively, i.e. using differential expression analysis. ‘‘Space” is a matrix which is built upon reference data. In this paper, reference data contains all Arabidopsis array samples that are measured by the Affymetrix Arabidopsis ATH1 Genome Array^3^. For each stress group in each data set, gene-wise fold changes are calculated between stressed plants and their corresponding controls. The fold changes fill one column of the space matrix. This generated matrix, which we call reference space (*S_r_*), contains all stress-relevant information represented in the reference data. In addition to *S_r_*, we calculated the mean reference space 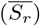. For constructing 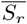, for each stress group, the gene-wise mean value of fold changes in *S_r_* is calculated and stored in a column in 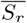. More detailed information, as well as a step-by-step guide for creating *S_r_* and 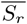, are provided in supplementary file 1.

##### Space Trimming

The stress grouping in this study is done based on the expert annotation provided alongside public data sets. Therefore, this grouping doesn’t necessarily reflect the different classes of biological mechanisms that shape the plant response spectrum. There are groups of biologically-related stresses, which in turn make some stress responses very similar in their full genome signature. Logically, stresses to which plants respond using the same common mechanisms and pathways, have similar gene expression fingerprints. On the other hand, stresses have significantly different gene expression patterns when few to no common genes are involved in their corresponding stress responses.

From the mathematical point of view, the distance between distinct stress responses manifests itself in the collinearity of axes of the extracted space. Collinearity in a mathematical space is a source of redundancy, and in our application, can result in lower accuracy and robustness.

We came up with a new algorithm named space trimming: an unsupervised approach which in combination with space extraction, makes up a hybrid method that can detect new groups of stress responses. We call these new-found groups meta-stress groups.

Space trimming uses a consecutive combination of hierarchical clustering and leave-one-out cross-validation (LOOCV) to remove the aforementioned redundancy from a space. Space trimming consists of three steps:

1. Clustering and cross-validation analyses are done on the space under study, and a dendrogram based on the calculated similarities is constructed.
2. Groups of stresses that are close and have low accuracy are combined to make meta-stress groups. Groups that merge under the 50% of the maximum height in the dendrogram (i.e. groups with the distance of 50% of the maximum distance or lower) are considered close, and groups of stresses that mostly have an accuracy of less than 0.7 are considered low in accuracy.
3. Any newly-generated meta group that has at least the same performance as its subgroups is kept. All other meta groups are reverted back to their former groups.

We applied the Space trimming algorithm to the reference space *S_r_*, generated in the last section, from Arabidopsis microarray data (Fig. 2).

**Figure 2:**
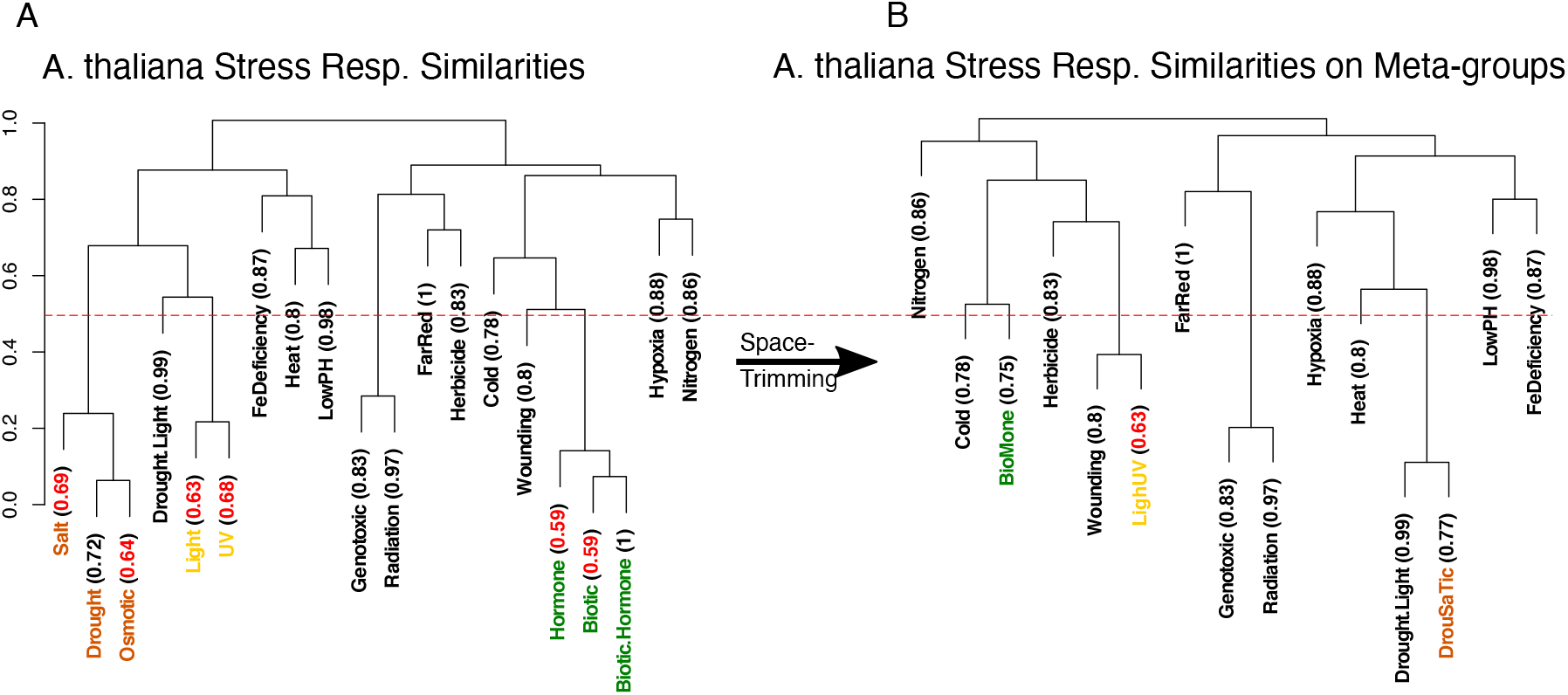
Space Trimming. Stress groups are clustered and for each group, leave-one-out cross-validation accuracy is calculated, written in parenthesis, as shown in panel A. Close groups with low accuracy, written in red, are combined to form new stress groups, called meta-groups, as shown in panel B. Groups are considered close if they merge in the dendrogram in a height lower than 0.5 (50% of maximum height). This cut-off height is shown in the figure with a dashed red line. In this figure, Salt, Drought and Osmotic stress groups, marked with brown color, merge into DrouSaTic meta-group, Hormone, Biotic and Biotic.Hormone groups form BioMone meta-group, written in green, and Light and UV groups combine into LighUV, shown in yellow.

In LOOCV, by definition, one sample is left out for testing. In our LOOCV scheme though, we left out one GSE (GEO^4^ series) data set in each cycle. Due to batch effects, even with proper preprocessing, samples from the same GSE set tend to be similar. With leave-one-GSE-out cross-validation, we make sure that the stress response from different data sets could be successfully matched together.

In each iteration, one GSE data set is chosen as the test set, it is mapped to the rest of the data sets, the training set, and it is counted as a successful match if the analyzed test data and its most similar data set from training set undergone the same stress group. Using the confusionMatrix function from the caret package [21] in R^5^, matching accuracy and robustness of the method is evaluated (Fig. 2A). With an overall balanced accuracy of 0.43, a Cohen’s kappa of 0.385, and an accuracy *p*-value of 7.35 × 10^−42^, PhysioSpace could successfully match the samples going through the same stress group.

As expected, clustering analysis exposed the similarities among different stress responses. For instance, responses to Osmotic, Drought and Salt stresses seem to have common underlying activated gene groups. Or regarding Biotic, Hormone, and Biotic.Hormone (double stress), their close proximity points toward a very similar stress response. They also predominantly have lower accuracy comparable to other stress groups (Fig. 2A). This led us to the assumption that these groups of stress responses share one or few underlying defensive mechanisms, such as an innate immune response.

Merging the similar stress groups and constructing the meta-stress groups result in an improved performance of the method (Fig. 2B). We constructed three new meta-stress groups: BioMone, which comprises of Biotic, Hormone, and Biotic.Hormone stress groups, DrouSaTic, that was built by combining Drought, Salt, and Osmotic stresses, and LighUV, which is made by merging Light and UV stresses (Fig. 2B). Redoing the LOOCV on the new grouped space demonstrates the performance gain, with an accuracy of 0.57 and a Cohen’s kappa of 0.49, which increased 0.14 and 0.105 respectively in comparison to the classical grouping of stresses. And the accuracy *p*-value stays significant, as it is equal to 2.91 × 10^−39^.

The resulted space, which we call meta-reference space or *S_mr_*, and its successive mean space which we denote by 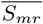, are the spaces we use as the reference throughout the result section, though it is possible to use *S_r_* and 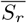 as a reference too, for example in cases that individual stresses are needed to be characterized, e.g. if Salt and Drought stresses are needed to be studied separately.

#### 2.2.2 Physio-mapping

After acquiring a space from known training data, we can map new data from any technology or species back into the space and find similarities between the new unknown information and the known training data. Physio-mapping is a nonlinear, model-free mapping, designed to take advantage of omics data structures and to compensate biases from heterogeneous assessment protocols. Omics are mostly framed in high-feature low-sample arrangements. In most cases, the majority of features in these types of data sets are intrinsically dominated by noise rather than physiologically informative facts about the samples under study. Results commonly acquired by differential expression analysis are great examples of this phenomenon; in most cases of differential expression analyses, only a small proportion of features in an omics data set can be found to be significantly different, i.e. correlated with circumstances that are being studied.

Presuming this assumption, the mapping is done by taking the following steps:

1. Either

a. A new space is extracted from the new data. This means that for each gene in each stress case, a fold change is calculated by modeling the gene behavior under the respective stress given the control. We call this new input space *S_i_*.
b. For each stress type *γ*, genes are sorted from the lowest to highest fold change.
c. *N* percent lowest and highest genes are selected as *L_L_*(*γ*) and *L_H_*(*γ*) for each *γ. N* is a user-defined parameter. In this paper, it is between 3 to 5 percent.

or Differential expression analysis is done on the input data, and for each stress type *γ*, down- and up-regulated gene sets are calculated, which are called *L_L_*(*γ*) and *L_H_*(*γ*), respectively.
2. For each axis on the reference space (i.e. each column in 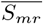), a statistical test is performed between *L_L_*(*γ*) and *L_H_*(*γ*) gene groups to form the *PS* matrix:

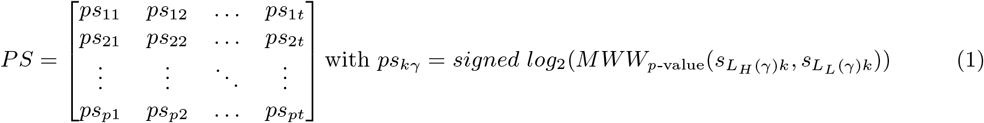 In equation 1, *ps_km_* is the physio score between the *m^th^* sample and the *k^th^* column of reference space 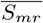. *ps_km_* physio score value shows how similar the *m^th^* sample of *S_i_* is to the *k^th^* column in 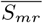. *MWW*_*p*-value_(*a,b*) is a function that calculates the *p*-value of a Mann–Whitney–Wilcoxon statistical test (also known as Mann–Whitney U test or Wilcoxon rank-sum test) between a and b. *s_L_H_(γ)k_* and *s_L_L_(γ)k_* are the sets of values in *k^th^* column, and *i* ∈ *L_H_* and *i* ∈ *L_L_* rows of 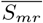, respectively. And *signed log*_2_(*x*) is 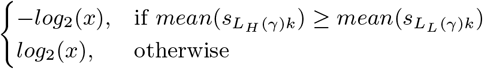

*PS* is a physio score matrix, containing similarity values of all input samples to all axes (i.e. columns) in the reference space. This physio score matrix is the output of Plant PhysioSpace, the scores quantify the intensity of different stress types in different samples, and they are comparable within the same physio score matrix^6^. To elaborate on how to interpret the physio scores, we analyzed the GSE13739^7^[22] data set from GEO using Plant PhysioSpace. In GSE13739, the responses of wild type and mutant *Arabidopsis thaliana* to *Golovinomyces orontii* (powdery mildew) are studied till 7 days post infection. The mutant in this study is Salicylic acid biosynthetic mutant *ics1*, which is expected to be more resilient compared to the wild type. We plotted the physio score matrix (calculated using equation1) as a heatmap (Fig. 3), with samples in columns and stress groups in rows of the heatmap, and the color showing the magnitude of each stress in each sample. Based on the experiment description, we anticipated 1) the plants to behave as they are under Biotic stress, and 2) the mutant plants to have a milder response compared to the wild type since they are more resilient. The results show similar dynamics in both wild type and mutant groups of samples: BioMone (Biotic or Hormone) is the dominant stress response, and also its corresponding score increases with time, with wild types reaching a stronger response levels compared to the mutants. Our results are aligned with what we expected from the experiment. In addition, they show that physio scores are comparable between samples (row-wise in the heatmap), as well as among stress groups (column-wise in the heatmap).

**Figure 3:**
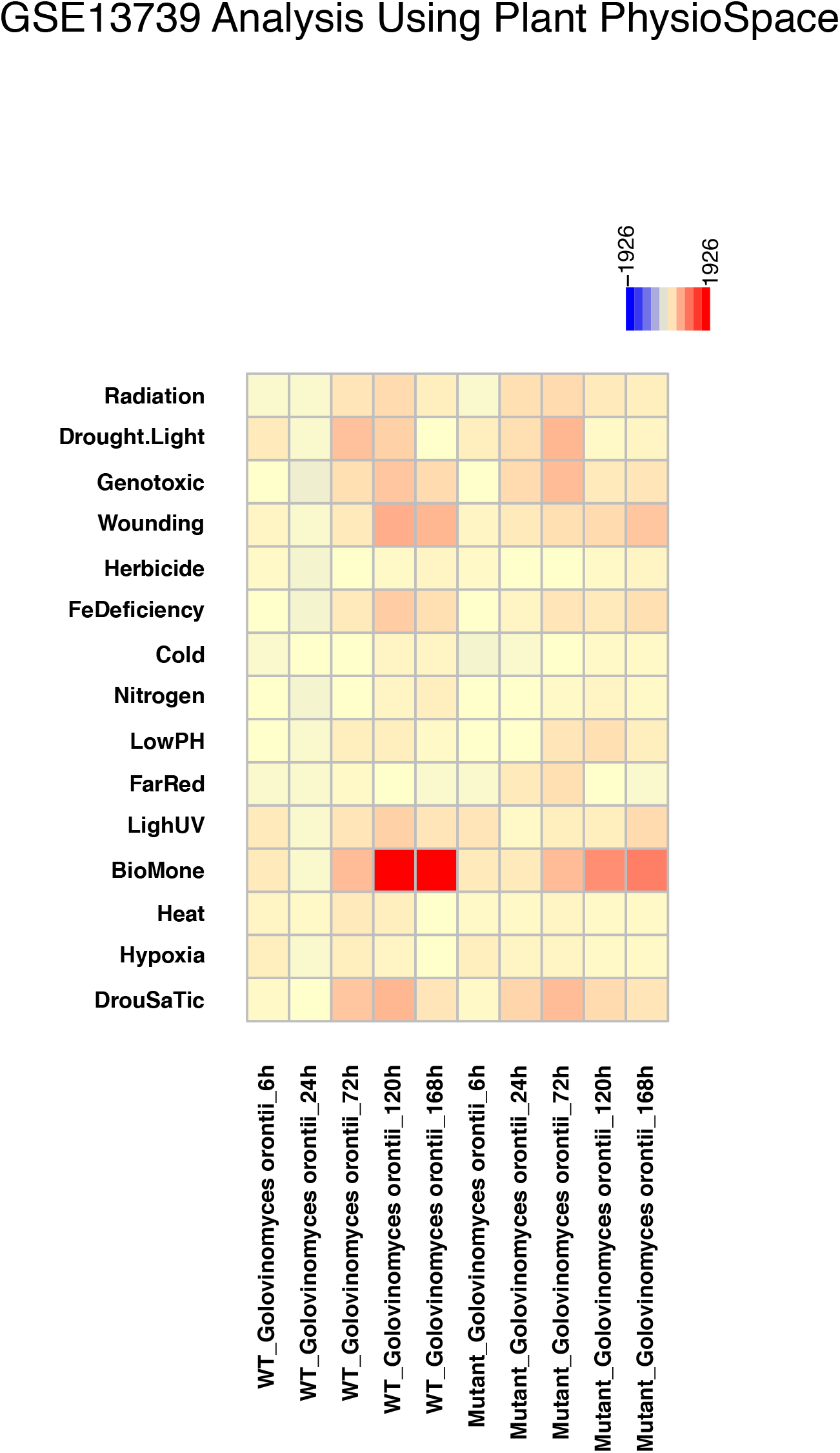
Analysis results of GSE13739. The GEO data set GSE13739 is mapped against the 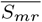 reference space using Plant PhysioSpace. The data set provides samples of Wild type (WT) and Mutant (MT) *Arabidopsis thaliana* plants that are infected with *Golovinomyces orontii* (Biotic stress). The mutant plants are expected to be more resilient.

For inter-species mapping, for instance in analyzing new data from *Oryza sativa* using a space generated from *A. thaliana*, we resorted to orthologous genes. By using the ideal assumption of orthologs to have identical biological roles in all species, we mapped genes to their orthologs in cases with interspecies translation. For the mapping, we utilized the Basic Local Alignment Search Tool BLAST [23]. More specifically, we used the BLASTn tool to align sequences from *Oryza sativa*, *Glycine max*, and *Triticum aestivum* against *A. thaliana*, with *α* = 10^−2^ as the cutoff for the “expect value”. Expect value, also known as expectation value or e-value, represents the number of random alignments with scores equivalent to or better than the resulted match [24]. In other words, the expect value depicts the statistical significance of the alignment. Therefore, in cases with multiple matching, the gene with the lowest expect value is chosen as the match, to have a one-to-one matching between genes from different species and *A. thaliana*.

## 3 Results

### 3.1 Stress Space Verification by GO Analysis

For substantiating the authenticity of stress information collected using our space generation process, we utilized the gene list analysis section of PANTHER [25, 26]. For each stress group in our generated mean meta-stress space 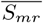, we selected the genes with an absolute score value of more than one and a half^8^, and tested this gene list by PANTHER overrepresentation Test, against Gene Ontology (GO) biological processes (Fig. 4 and supplement files 2 and 3). From 15 different stress groups, 11 were found to have the GO terms corresponding to the same stress enriched, with significant corrected *p*-values of less than 0.001.

**Figure 4:**
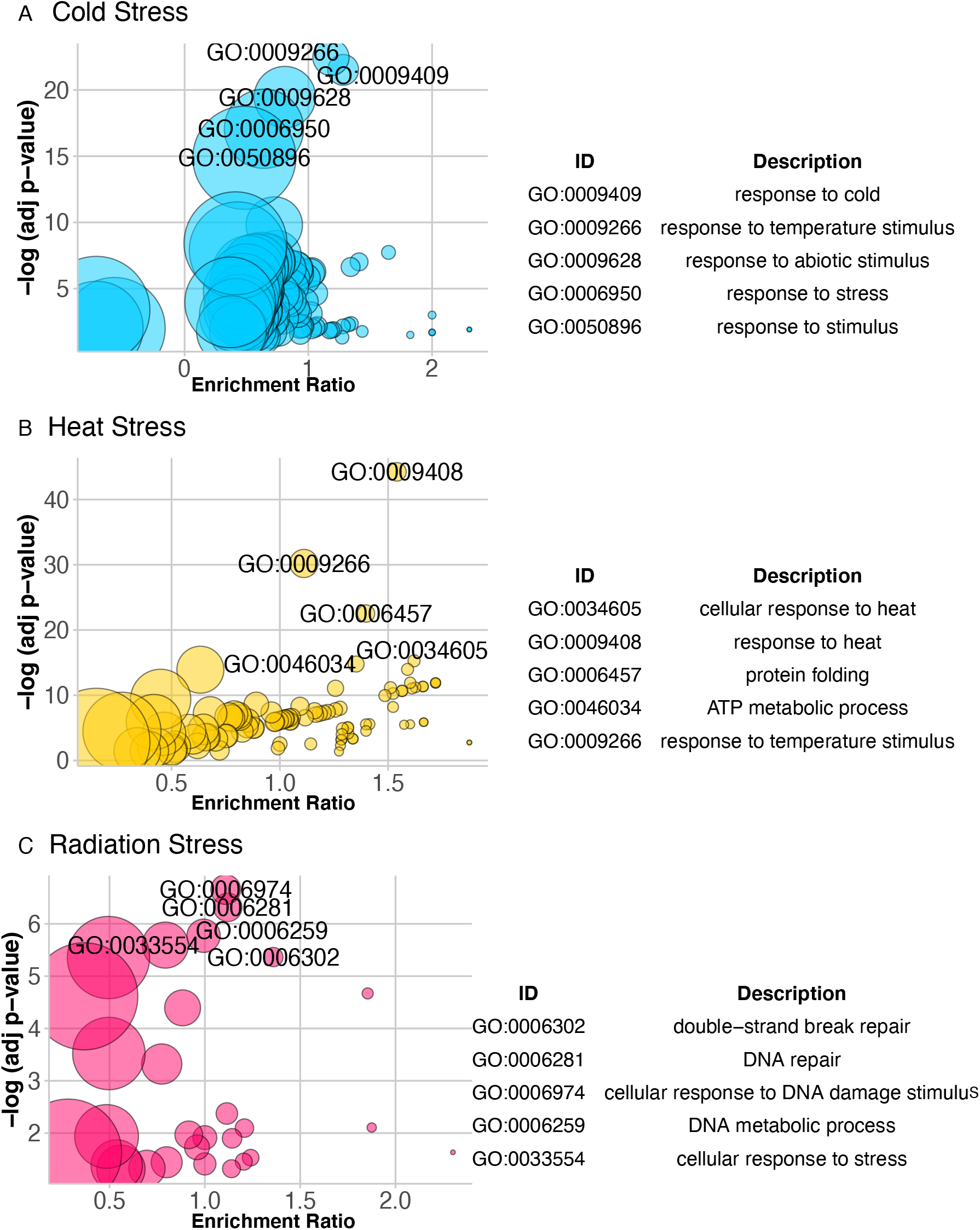
GO Analysis of Mean Stress Space. Results of GO analysis on three stress groups are demonstrated using bubble plots. In the plots, each enriched GO term is represented by a circle, with adjusted *p*-values as y-axis and enrichment ratio as x-axis. The size of the circle shows the size of the gene list of the corresponding GO term. And enrichment ratio here means the ratio between the actual number of differentially expressed genes and the expected, in each GO group. For each plot, 5 most significant GO terms are labeled on the plot and listed in a table beside each plot. Complete set of bubble plot and set of significant GO terms for all 15 stress groups are provided in supplement files 2 and 3. Plots were generated using the GOplot package in R [42].

### 3.2 Inter-Technology Translation

Next generation sequencing (NGS) has revolutionized the biological sciences. Its speed, cost and data quality outpaced the older DNA-microarray technology, which is why NGS became the standard method to study transcriptomes. Yet, microarrays were used for RNA quantification for decades. The vast microarray backlogs have the potential to grant an invaluable resource for new biological studies. Unfortunately, the measurement technology has an inevitable impact on the transcript measurement levels and the distribution of the resulting data.

Data derived from different platforms are distinctly different. Hence, there are numerous methods to translate measurements from one technology to another [27, 28]. Moreover, with the third generation sequencing right around the corner (PacBio and Nanopore, to name a few [29]), there is a high demand for computational methods capable of transferring useful information between different measurement technologies.

Since PhysioSpace utilizes the differential expression relations of the genes and not absolute values for space generation and mapping, it can translate between each and any technology, as long as there exists a proper method for detecting the differentially expressed genes in the mentioned technology. As proof for this claim, we mapped more than 900 RNA-seq samples into the microarray space *S_mr_* (Table 1). Our method can map the same stress type from microarray to RNA-seq data with 78 percent accuracy. We also calculated the probability of acquiring this accuracy by chance, by randomly permuting the sample labels and calculating the random accuracy. The performance of our method is significantly higher than any random accuracy we acquired^9^, with a *p*-value of less than 10^−7^.

**Table 1:**
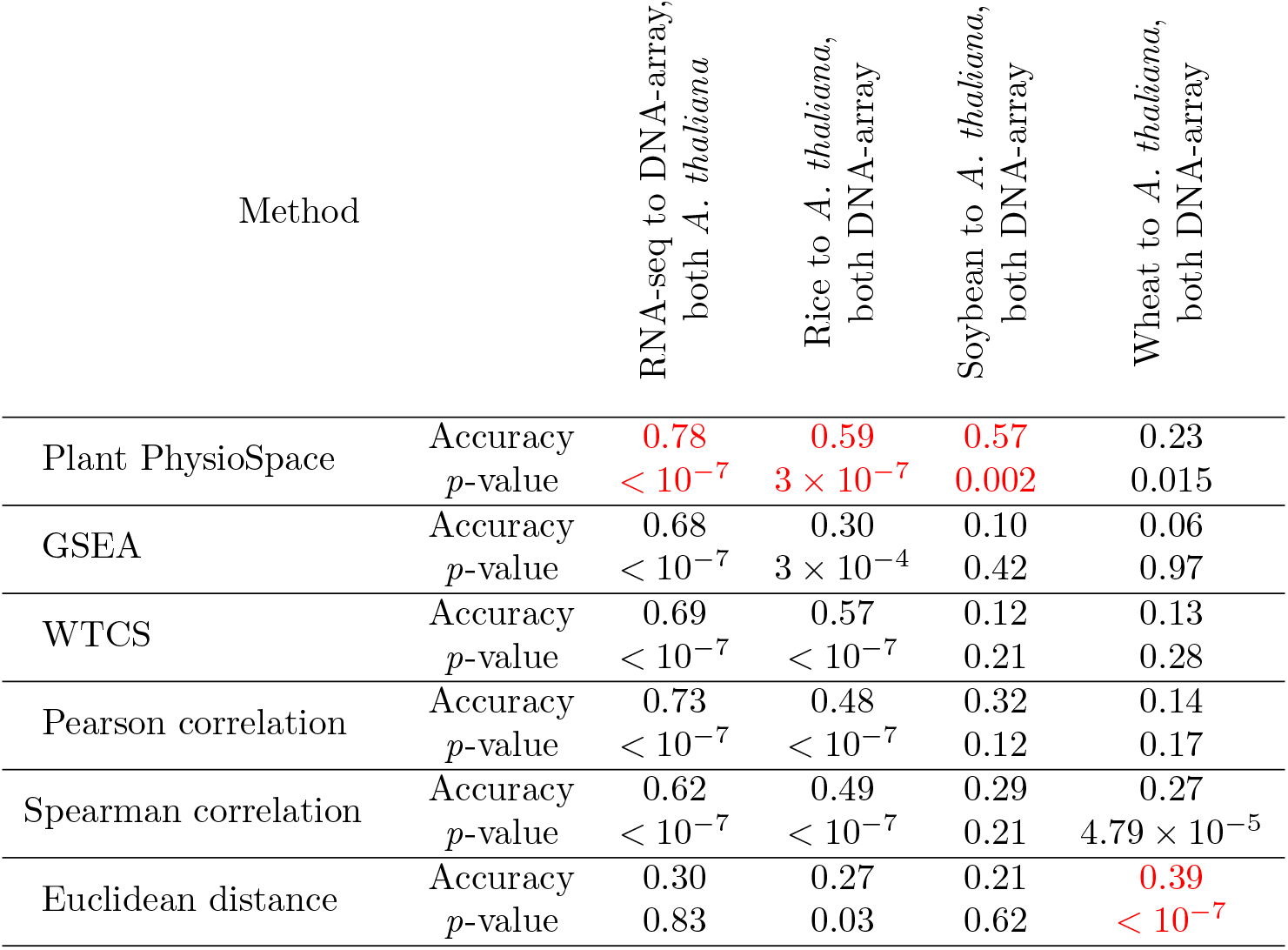
Stress translation between platforms and Species. In each column, the best performer is marked in red.

### 3.3 Inter-Species Translation

Although not agriculturally relevant, Arabidopsis is arguably the most investigated species in plant sciences. Its availability, compact size, and fast growth made it an ideal model species. Nevertheless, there are significant differences between the Arabidopsis plant model and crop plants, necessitating procedures for converting well-studied physiological knowledge, e.g. regarding plant response to different types of stress, from Arabidopsis to crops. In this section, we show how Plant PhysioSpace can be utilized for this purpose.

We chose three of the most commercially relevant crops to study: *Oryza sativa* (rice), *Glycine max* (soybean) and *Triticum aestivum* (wheat). For each crop, more than 100 microarray samples of stress response experiments were curated, normalized, preprocessed and mapped to the Arabidopsis space *S_mr_*. For *Oryza sativa* and *Glycine max*, Plant PhysioSpace achieved respective accuracies of 59 and 57 percent, both of which were significantly higher than any accuracy earned by chance. On the other hand, translation of stress response from *Triticum aestivum* DNA array data to *A. thaliana*, with an accuracy of 23 percent and a *p*-value of 0.015, was not successful (Table 1). In section 3.5 of this paper, we provided a thorough investigation into wheat to Arabidopsis translation, hypothesized and examined the reason behind the translation failure, and provided solutions for fixing it.

### 3.4 Benchmarking Plant PhysioSpace against Other Methods

We used the results from inter-technology and -species stress response translation to benchmark our method. Plant PhysioSpace is compared to the most common approaches used in bioinformatics for measuring relations between two or more gene expression samples: Gene Set Enrichment Analysis (GSEA) [30], Weighted Connectivity Score (WTCS), which is an advanced version of GSEA used in connectivity map [31], Pearson and Spearman correlations, and Euclidean distance. For each method, fold change values of samples, from different technologies and species, are calculated and used for finding the similarities between samples. Based on our results in inter-species and -platform mapping, Plant PhysioSpace could outperform other methods in all scenarios, except in mapping from wheat to Arabidopsis (Table 1).

### 3.5 In-depth Investigation of Wheat Stress Response

The poor performance of our method in translating stress response from *Triticum aestivum* to *A. thaliana* may have potentially derived from the microarray used to measure the wheat gene expressions. All microarray samples of *Triticum aestivum* in this study are generated by using Affymetrix Wheat Genome Array^10^. Not only the aged technology could potentially deter the accuracy of transcription measurements, but also as a polyploid, the complex genetics of wheat would make the task of measuring its RNA levels troublesome. Moreover, among all translations tested, wheat microarray provides the least amount of mapped orthologs: when translating from RNA-seq to microarray, we could find 12146 common features (i.e. genes). This amount was lower for both translations of rice and soybean to Arabidopsis, as the ortholog mapping could find 9455 and 9452 common features for them, respectively. And, in translation from wheat to Arabidopsis, 6202 common features could be found using the ortholog mapping, which is the least among the four translations. These common feature counts are correlated with the translation performance, considering the RNA-seq to DNA-microarray has the highest and wheat to Arabidopsis has the lowest accuracy, while rice and soybean mappings are somewhere in between (Table 1). This correlation suggests one possible reason behind the failed wheat translation is the low amount of orthologs. Fortunately, advances in NGS gave rise to wheat stress response data sets with higher precision and a wider range of sequence reads. In this section, we repeated the wheat-to-Arabidopsis translation from inter-species analysis, except with wheat RNA-seq instead of microarray data.

We turned to the Wheat Expression Browser [32, 33] as the source of *Triticum aestivum* RNA-seq data. From this source, we queried all data sets which study stress response, contain more than 30 samples, and include controls. Upon the first inspection, 8813 orthologs could be found between these data sets and the genes available in our Arabidopsis reference space, which in addition with the improved data quality provided by RNA-seq, increases the prospect of Plant PhysioSpace correctly detecting and quantifying the stress in these data sets. We mapped these data sets into mean meta-reference space 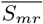, and plotted Physio scores of three stress groups with the highest values (Fig. 5).

**Figure 5:**
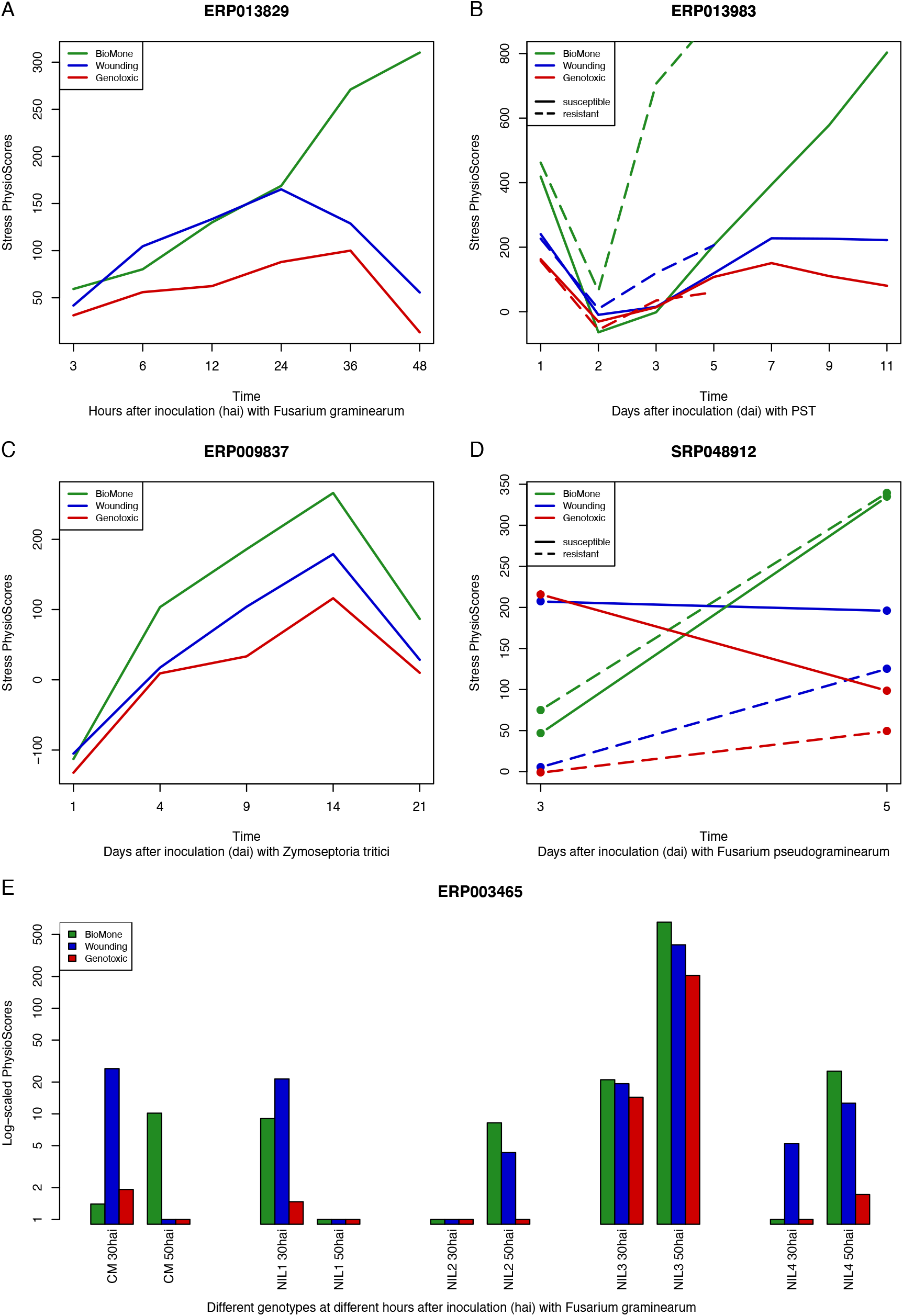
Time Series Analysis of Biotic Stress Response of Wheat RNA-seq data. 5 different biotic-stressed data sets from Wheat Expression Browser are mapped to the Arabidopsis space 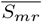, and the three groups with highest stress values are plotted for each data set. In 4 out of 5 cases, BioMone (Biotic and Hormone) stress group has the highest similarity value, with resistant mutants having higher responses than the susceptible ones (panels A, B, C, and D).

In the experiment set ERP013829, wheat response after inoculation with fungal pathogen *Fusarium graminearum* is measured through time [34]. For this experiment, Plant PhysioSpace correctly predicts that wheat is experiencing BioMone (Biotic and Hormone) stress. In addition, the diachronic rise in the response indicates how Physio scores are quantitatively comparable (Fig. 5A).

In the data set ERP013983, responses of two different mutants of wheat are studied to wheat yellow rust pathogen *Puccinia striiformis f.sp. tritici* (PST). The authors focused on the pathogen suppression of basal defense in plants [35]. From their results, they deduced the pathogen overcame the defense by rapidly suppressing the genes involved in chitin perception on day 2 after inoculation. In the susceptible interaction, this provides the possibility of invasion and colonization, while in resistant plants, this suppression is quickly reverted. Plant PhysioSpace results expose this mechanism correctly: Both plant types are inducing a BioMone response on day 1, followed by a suppression of the plant defense response on day 2. Eventually, the quick resurgence of intense BioMone response in resistant wheat helps it in withstanding PST, while the reaction of the susceptible trait might be too slow to deflect the pathogen (Fig. 5B).

Plants in ERP009837 went through the infection cycle of the hemibiotrophic fungus *Zymoseptoria tritici*. Similar to ERP013829, plants respond to the pathogen with a dominant BioMone response (Fig. 5C). Although unlike ERP013829, in which the experiment spanned a few hours, in ERP009837 plants were studied for a longer period. Physio scores suggest that wheat responds to the presence of the pathogen by increasing its BioMone response. This response starts to degrade from day 14, which is in alignment with the original publication of the data set [36], in which the authors state that on day 21, plant tissue is completely defeated.

In SRP048912, the responses of two different traits of resistant and susceptible wheat to Fusarium crown rot are studied [37]. Only two different time points are included in this experiment: 3 and 5 days after inoculation (dai). Plant PhysioSpace results correctly suggest the most dominant stress response present in plants is BioMone, with the resistant wheat having a stronger response than the susceptible (Fig. 5D).

Among the Wheat data sets we analyzed in this section, ERP003465 is arguably the most complex, and consequently most interesting as a testing scenario for our method. ERP003465 examined the behavior of 5 different genotypes under the disease pressure of *Fusarium graminearum* [38]. Two well-validated and highly reproducible QTLs (quantitative trait loci), *Fhb1* and *Qfhs.ifa-5A*, are studied from samples taken 30 and 50 hours after inoculation (hai). Five different genotypes were investigated: CM-82036, a progeny of the resistant Sumai-3, and four near-isogenic lines (NILs) bearing either, both, or none of the resistant alleles *Fhb1* and *Qfhs*. Among the four, NIL1 is a mutant with both QTLs, expected to have the highest resistance after CM-82036, NIL2 and NIL3 are mutants harboring *Fhb1* and *Qfhs* QTLs respectively, with both predicted to behave moderately resistant, and NIL4 missing both QTLs, and is likely to be susceptible.

Data analysis in the original paper was mainly based on differential expression analysis. As a first step, the total number of differentially expressed genes for each genotype at each time point was taken as a surrogate for stress response intensity to *Fusarium graminearum*. In the next steps, the weighted gene co-expression network analysis (WGCNA) was used to detect clusters of genes with similar patterns, and Gene Ontology analysis was utilized to infer the role of each cluster in the stress response.

Being able to quantify the intensity of each stress type at each time point, Plant PhysioSpace can provide much more insight into the characteristics and dynamics of the stress responses that are at play in the ERP003465 experiment (Fig. 5E). As this data set encompasses a high number of samples distributed between only two time points, we plotted the results as a bar graph. And because the results cover a wide range of values for this experiment, we used log-scaled Physio scores in the graph, and replaced values smaller than one by one (i.e. zero in log-scale).

Among the concluding remarks in the original paper, some are in concordance with the results from our method. For example, lines lacking *Qfhs.ifa-5A* are regarded as “slow responders” by the original authors, since they lack resistance against initial infection inferred by *Qfhs.ifa-5A*. This lack of early response can be seen in our results (Fig. 5E): lines lacking *Qfhs.ifa-5A*, i.e. NIL2 and NIL4, have no BioMone (Biotic and Hormone) stress response at the early time point, while NIL1 and NIL3 show a considerable BioMone stress response at the same time point. Another remark from the original paper suggested that a lack of timely defensive reaction could result in a higher infection in a later time, and consequently stronger response, and vice versa: a quick response may reduce the intensity and infection at a later time. This can be seen in the contrasting response dynamics of NIL1 versus the other lines (Fig. 5E). NIL1 possesses both QTLs: *Qfhs.ifa-5A* ensures an early and fast stress response, evident on 30 hai time point. And a strong follow up, courtesy of *Fhb1*, results in a non-existent BioMone response at 50 hai. NIL3 contains *Qfhs.ifa-5A*, so it benefits from a quick response at 30 hai, but due to the absence of *Fhb1*, it cannot be rid of infection at 50 hai, evident by the high BioMone response at that time point. As mentioned, lines NIL2 and NIL4, which lack *Qfhs.ifa-5A*, do not have an early response and have to play catch up with other lines on the later time point.

Although many conclusions that could be derived from our method are similar to the ones from the original publication, there are some discrepancies between the two groups as well. For instance, in most samples, Wounding stress response is not only present, but it is even stronger than BioMone response in some cases. This is in contrast with the original paper, in which it is mentioned that inoculation was done cautiously without wounding the tissue. Interpretation of CM-82036 defensive behavior is another point of difference between our method and the results from the original paper. Kugler et al. construed the high number of differentially expressed genes (DEG) at 30 hai as a sign of strong early response for CM-82036, even stronger than NIL1 and NIL3. They followed up by studying specific gene families that are relevant to defense mechanisms, such as UGTs and WRKYs, and showed more DEGs from these families can be found at 30 hai in CM-82036 versus other lines. This finding is different from what we can interpret using our method: Although CM-82036 exhibits BioMone response at 30 hai, the magnitude is somewhere between fast responder lines, that is NIL1 and NIL3, and slow responders, i.e. NIL2 and NIL4.

We speculate the main reason for the aforementioned inconsistencies is the particular way the preprocessing was done in the original paper. In their preprocessing, Kugler et al. mapped the reads to a list of barley high confidence genes and only used the reads with a possible match. This step drastically reduces the number of analyzed transcripts, and also discards wheat-specific genes with no barley homologs. Our method is designed for high-dimensional data, preferably data from the whole genome, therefore the specific preprocessing of this data set might have reduced the performance of Plant PhysioSpace. We should also mention that stress responses are not mutually exclusive; A plant can display multiple different responses at the same time, some of which may even share part of their biological pathways. *Fusarium graminearum* could have damaged the plant tissue at some point, which explains the existence of wounding response alongside BioMone.

Albeit the mediocre results of the last experiment, in this section we showed how, in 4 out of 5 data sets, Plant PhysioSpace could:

1. correctly identify the type of stress plants are going through.
2. accurately relate the response from RNA-seq test data to DNA-array trained models.
3. rightly translate *T. aestivum* stress response to *A. thaliana*.

### 3.6 Plant PhysioSpace Application in Single Cell Analysis

Single cell technologies facilitate investigating transcription profiles in single cell resolution, in order to perceive the genetic basis of each cell type and its function. Although relatively new, more and more plant single cell data sets are becoming available to the community [9, 10, 11, 12, 13]. For now, most sequence data sets are focused on Arabidopsis roots. They try to gain an in-depth understanding of transcription patterns of different cells in different developmental stages of wild-type non-stressed plant roots. To our knowledge, the only publication in which stressed single cells were sequenced is the paper by Jean-Baptiste et al. [12]. In this work, 38° C heat stress was applied to 8-day-old seedlings for 45 minutes. Subsequently, roots of the seedlings were harvested, along with the roots of age- and time-matched control seedlings. The authors could capture and sequence 1,009 cells from the stressed group and 1076 from the control group. For processing the sequencing results, they followed the usual single cell analysis pipeline: PCA, UMAP and clustering, followed by differential gene expression analysis on clusters and enrichment tests on genes related to heat-shock. The results show the “promise and challenges inherent in comparing single cell data across different conditions and treatments”. In this section, we demonstrate how a dedicated method, such as Plant PhysioSpace, can bring forth more benefits than using the methodological norms.

To analyze the single cell data set, we used the gene-wise mean value of all control cells as the reference, calculated fold changes for each single cell, and fed those fold change values into the Plant PhysioSpace pipeline (Fig. 6). Regardless of the cell type, heat-stressed single cells had significantly higher heat stress scores, compared to control single cells (Fig. 6A). For studying the heat-induced cell type disparity, we overlaid heat stress scores on UMAP and tSNE plots (Fig. 6B&6C). In both tSNE and UMAP plots, coordinate values calculated in the original paper of Jean-Baptiste et al. were used. As a result, cells are bundled in cell type clusters in the UMAP plot, while in the tSNE plot, cells are clearly separated into two big clusters of control and stressed. Although, inside these two big clusters, sub-clusters of different cell types are evident (Fig. S1). On the UMAP plot on the other hand, big clusters represent cell types (Fig. S2), and inside each cell type cluster, groups of control and stressed cells may or may not be distinct, depending on the cell type. For example, in Hair and Non-hair clusters, control and heated cells are separated, while the separation is less pronounced in Stele cells (Fig. S2).

**Figure 6:**
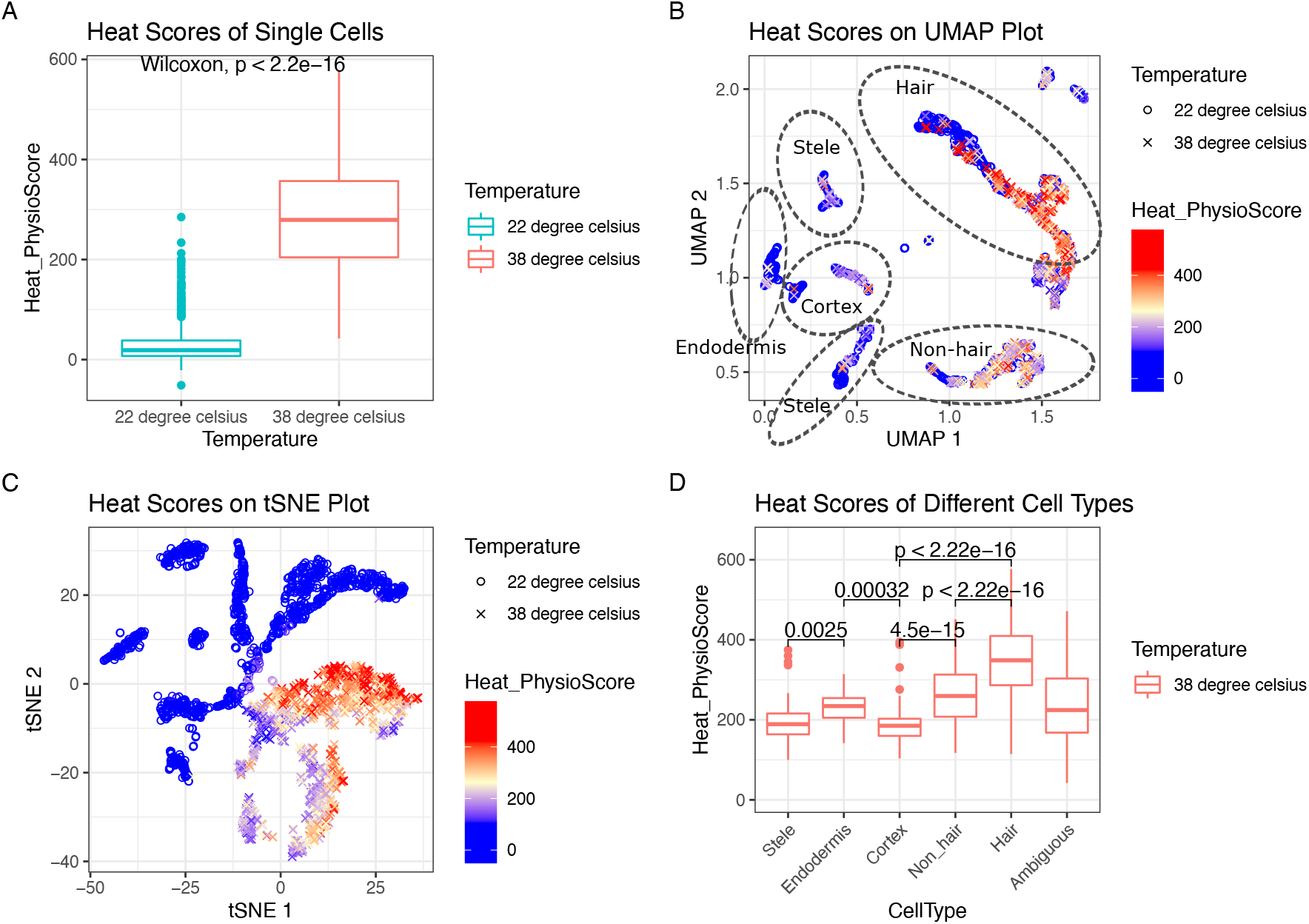
Single Cell Analysis Results of Plant PhysioSpace. Stress scores were calculated for each cell. For demonstrating the outcome, we plotted the heat score of the two big groups of control and stressed, shown in panel A. This box plot proves how Plant PhysioSpace could correctly detect and quantify stress response in single cell data. On panels B and C, we overlaid the heat scores on UMAP and tSNE plots, respectively. On panel D, boxplot of heat scores, on y-axis, were plotted against different cell types, on x-axis. Cell types on the x-axis are ordered based on the morphological anatomy, starting from inner cell types to outermost cell layers (excluding Ambiguous cells, which come at the end).

To look into the distinct behavior of different cells under stress, we also plotted cell heat scores grouped by the corresponding cell types (Fig. 6D&S3). The results show how Hair and Non-hair cells have higher heat scores, which demonstrates how the outer layers of roots are sharper in their response to heat. This finding is in concordance with one of the conclusions in the original paper, in which based on the behavior of the heat-relevant genes, they concluded the three outermost cell layers of the root went through higher levels of changes caused by the heat stress. The authors hypothesized this may be because of more direct exposure of the outer layers to the heat shock, resulting in a quicker and stronger response.

Although resulting in generally the same conclusions, in this analysis Plant PhysioSpace provided an advantageous experience for the end-user, through providing:

1. convenience: unlike the original paper, there was no need for search and curation of heat stress gene clusters, as they are already available in Plant PhysioSpace, as well as clusters for other common stresses.
2. precision: not only the stress type but also the magnitude of the stress response could be quantified by our method, something which is lacking in traditional gene list enrichment approaches. For example, Plant PhysioSpace results suggest a stronger response in Hair response, compared to Non-hair response (Fig. 6D). This inference could not be concluded by the results of traditional methods.
3. optimization: in one run, our tool calculated responses of 20 different stresses for 2085 single cells, in less than 3 minutes on a 2-core laptop CPU. This swift performance is accomplished by precalculating the stress space, in combination with an optimized mapping algorithm, all of which is readily available for the community to use.

### 3.7 Availability

To provide the community with an easy-to-use implementation of our method, we built Plant PhysioSpace into two different R packages: a method package (https://git.rwth-aachen.de/jrc-combine/PhysioSpaceMethods) containing functions for generating new spaces and Physio-mapping, and a data package (https://git.rwth-aachen.de/jrc-combine/PlantPhysioSpace) comprising plant stress spaces such as 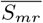 and 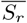 that were used in this paper.

In addition, we made a shiny^11^ web application of Plant PhysioSpace (Fig. 7). We hosted the web app on shinyapp.io (http://physiospace.shinyapps.io/plant/), to be freely available to use (under the terms of GPL-3 license). We also built a Docker image of the ready-to-use tool (https://github.com/usadellab/physiospace_shiny).

**Figure 7:**
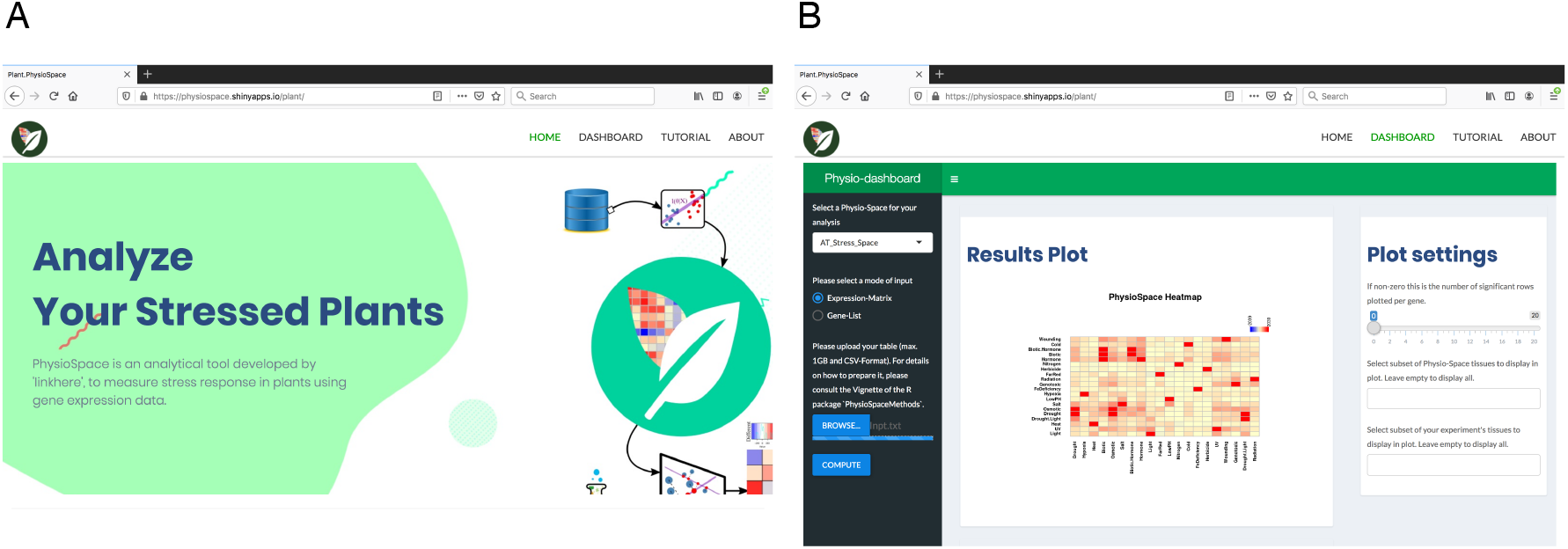
Plant PhysioSpace Web-application

### 3.8 Supplemental Material

This paper is accompanied by three supplement files and three supplement figures:

1. **Supplementary file 1** is a PDF text file that provides a detailed explanation about how “space extraction” is performed.
2. **Supplementary file 2** is a compendium of all 15 bubble plots related to section 3.1, “Stress Space Verification by GO Analysis”.
3. **Supplementary file 3** is an excel file that contains the results of the PANTHER GO analysis for all the 15 stress groups. In the file, correctly detected GO terms are highlighted, and stress groups that had irrelevant GO terms enriched are labeled.
4. **Supplementary figures** are related to section 3.6, “Plant PhysioSpace Application in Single Cell Analysis”. Supp. figures S1 and S2 are alternative versions of figures 6C and 6B, in which cell types are annotated in place of temperature. And supp. figures S3 is the same as figures 6D, except it contains the control cells physio scores as well.

## 4 Discussion

Gaining proper insight into stress response mechanisms in plants is not only a must for the future of agricultural research, but will prompt advances in the plant research field in general. In this study, we developed an advanced computational method, designed to aid in understanding stress response in plants. The lightweight algorithm allows it to run on either personal computers, or as a web application, making it an ideal tool for experimental quality control, data set annotation, to draw conclusions considering thousands of genes, et cetera.

We built the new method upon a previously published method in humans, called PhysioSpace. We achieved this conformity by curating a multitude of Arabidopsis stress response samples to have a rich training data set, adapting the space generation algorithm, i.e. training, to acclimate to the specific characteristics of stress response data in plants, and thoroughly testing against other species and types of data. The results of this study demonstrated that Plant PhysioSpace can be a convenient and practical tool for analyzing new stress response data sets, to apprehend, contrast to state of the art, or to simply quality control.

Notably, our tool could perform adequately even when it was mapping information between different platforms and species. Although, it is crucial to bear in mind these cross translations necessitate for some conditions to be true. In cross-platform translation, it was assumed that with the same experimental setup and samples, there are computational pipelines available which roughly compute the same differentially expressed gene lists regardless of the platform used. And in cross-species mapping, we assumed orthologous genes have the same biological function across all species; evidently impossible to be consistently true for all genes, but a sizable portion of genes have to pass this criterion for the inter-species translation to work.

We demonstrated how Plant Physiospace can provide insights when used for analyzing single cell data sets. Recent advances in single cell technology call for suitable bioinformatic analysis tools, for example for reducing the interfering technical noise [16]. The clear, factual results derived from single cell data analysis in this paper bring a spectrum of applications to mind for the future, especially in the light of approaching plant single cell atlas projects [39, 40].

Plant PhysioSpace may seem paradoxical at first, as it is explained to be a ‘dimension reduction” method that extracts stress information from gene expression data ‘without reducing dimensions”. Yet, this ostensible contradiction is plausible in actuality, since our method doesn’t discard any features from the training data when extracting stress information in the space generation stage (i.e. model training), but only removes the irrelevant features in the physio-mapping (i.e. when applying the method to the new samples). As mentioned before, the majority of genes remain unaltered under stress. We showed this unvarying behavior using the mean meta-reference space 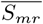(Fig. 8). Taking the absolute fold change value of 1 as a cutoff, only a small portion of plant genes are changing under stress. To be more specific, from all 22249 genes available in 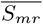, only 2905 (~13%) change at least under one stress type. In addition, the number of changed genes is a function of the stress type, ranging from ~0.07% in BioMone (biotic, hormone, or both stresses) to ~5.8% in the Drought.Light (double stress) group (Fig. 8A). Moreover, the majority of varying genes are specific to a stress group, as 2175 out of the 2905 genes (~75%) only change in one stress group (Fig. 8B). The low dimensionality of the stress responses, along with the highly specific features, may suggest that one can remove the genes with the absolute fold change of less than 1 in the training step, as it seems that keeping all the features till the end is not beneficial, while removing them could result in an even more compressed model. We are against this exclusion, however, on the ground that the removal of genes with an absolute fold change of less than 1 can eliminate relevant information. As proof, we ran the single cell processing of the last section once more, but this time as a reference, we used a “reduced” 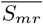, which is a version of 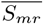 with all the 2905 varying genes removed (Fig. 8C). Although not as high as their counterparts calculated by using the complete 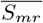, heat scores calculated using the reduced space are still significantly higher in heated single cells compared to the control cells (Wilcoxon rank-sum test *p*-value < 2.2 × 10^−16^), showing that the less varying genes also carry stress-related information. Therefore, we conclude the benefits of keeping all possible features outweigh the advantages that come about by removing them. Having the information from the whole genome at hand is especially helpful when analyzing data sets that don’t include a high number of genes, because the overlap between the features present in the model and the new data may become too small. Many new sequencing technologies, the Oxford Nanopore MinION [41] for example, or novel single cell sequencing platforms, provide readings on a limited number of genes, and keeping the whole genome in the trained model can boost the performance of our method on data from the aforementioned technologies. That being said, a proper and clever way of reducing dimensions of the trained spaces, a far more complex way compared to the fold-change cutoff scenario we examined before, has the potential to increase the efficiency and reduce the complexity of our method.

**Figure 8:**
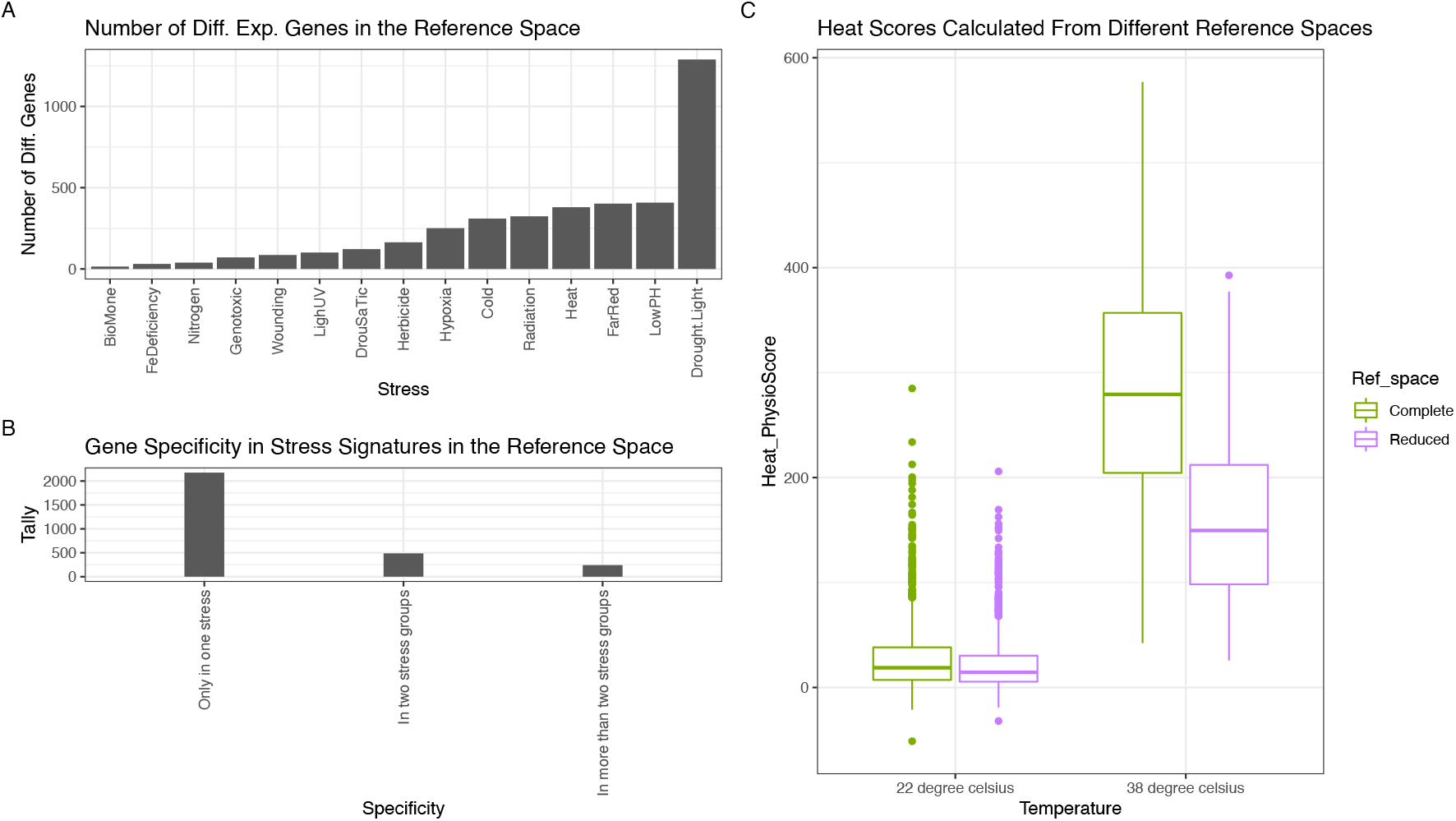
Differentially Expressed Genes in the Reference Space. Behavior of genes with an absolute fold change value of more than 1 in the reference space 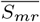, and their effects on the performance of 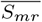 as a reference space is examined. In panel A, the number of genes with an absolute fold change value of more than 1 in each stress group is demonstrated. Since there are 22249 genes available in the space 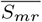, the ratio of differentially expressed genes to all genes among different stress groups spans from around 0.07% in BioMone (biotic, hormone, or both stresses) to around 5.8% in the Drought.Light (double stress) group. In panel B, we explored the specificity of differentially expressed genes to stress groups. Among all 22249 genes in 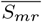, only 2905 (~13%) are differentially expressed in one or more stress groups. From these 2905 genes, 2175 (~10%) are specifically expressed in only one stress group, 488 (~2%) are expressed in two stress groups, and 242 (~1%) in more than two stress groups. Hence, we conclude that the majority of expressed genes in the reference space 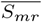 are specific to one stress group. In panel C, we used the heated single cells to study the effect of the 2905 differentially expressed genes on applicability of 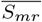 as a reference space. We compared the performance of 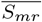 against 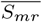 without the 2905 differentially expressed genes, which we called the ‘‘reduced” 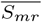. As evident from the boxplot, the heat scores are still significantly different between control and heated cells, even when the reduced space is used, although the magnitude of the heat scores is decreased compared to when the complete space is used.

To our knowledge, Plant PhysioSpace is the only computational tool available capable of quantitizing stress response in plant cells. Therefore, it can be used to assess each cell under stress, to grasp an understanding of the complex responses and interplay of cells in plants under stress, and to achieve a comprehensive characterization of plant response to stress as a whole.

## 5 Data Availability Statement

All the scripts that generate the results of this paper can be found in https://git.rwth-aachen.de/jrc-combine/PlantPhysioSpacePaper.

## Footnotes

1 https://www.ncbi.nlm.nih.gov/geo/

2 https://www.ncbi.nlm.nih.gov/sra/

3 https://www.ncbi.nlm.nih.gov/geo/query/acc.cgi?acc=GPL198

4 https://www.ncbi.nlm.nih.gov/geo/

5 https://www.r-project.org/

6 The possible range of the physio scores is dependent on the number of shared features (i.e genes) between the test data and the reference space. Therefore, if in two different mappings, the numbers of shared features are the same, the scores from the two different resulted physio score matrices are comparable as well.

7 https://www.ncbi.nlm.nih.gov/geo/query/acc.cgi?acc=GSE13739

8 For BioMone and FeDeficiency stresses with cutoff of 1.5, less than 10 genes were selected, which is too small of a set for list enrichment analysis. Hence in these two cases, cutoff is reduced to one.

9 The highest acquired accuracy from 10,000,000 random runs for RNA2DNA translation was 52% (minimum = 12.08%, first quartile = 28.12%, median = 30.87%, third quartile = 33.56% and maximum = 52.35%).

10 https://www.ncbi.nlm.nih.gov/geo/query/acc.cgi?acc=GPL3802

11 http://shiny.rstudio.com

