## Supplemental File 1 for "Plant PhysioSpace: a robust tool to compare stress response across plant species"

### Supplement File 1: Space Extraction in Detail

Esfahani et al.

As mentioned, in PhysioSpace method, all samples are analyzed contrastively, i.e. using differentially expression analysis. We assume

$$X = \begin{bmatrix} x_{11}^{g_1} & x_{12}^{g_1} & \dots & x_{1m}^{g_p} \\ x_{21}^{g_1} & x_{22}^{g_1} & \dots & x_{2m}^{g_p} \\ \vdots & \vdots & \ddots & \vdots \\ x_{n1}^{g_1} & x_{n2}^{g_1} & \dots & x_{nm}^{g_p} \end{bmatrix} \tag{1}$$

is the $m$ by $n$ expression matrix from which we want to build a space, with $x_{ij}^{g}$ being the expression of $i^{th}$ gene in the $j^{th}$ sample. $j^{th}$ sample belongs to group $g$, and $g$ is from a set of control and all the stress labels from the stress set defined in data preparation section ($g \in \{Control, Biotic, Cold, Drought, \dots\}$). We also assume $n >> m > g$, which means multiple columns can belong to the same stress group $g$. Each row in $X$, i.e. each gene, is modeled separately:

$$x_i^s = f_i(x_i^c), \text{for each gene i} \tag{2}$$

In the equation 2, $x_i^s$ and $x_i^c$ are expression values of $i^{th}$ gene in stressed and controlled groups, respectively. The model $f$ is calculated using limma package [1] for microarray samples and DESeq2 [2] for RNA-seq data sets. Based on $f$, for each gene in each stress case, a fold change is calculated and stored in a space matrix:

$$S_r = \begin{bmatrix} s_{11}^{g_1} & s_{12}^{g_2} & \dots & s_{1p}^{g_p} \\ s_{21}^{g_1} & s_{22}^{g_2} & \dots & s_{2p}^{g_p} \\ \vdots & \vdots & \ddots & \vdots \\ s_{n1}^{g_1} & s_{n2}^{g_2} & \dots & s_{np}^{g_p} \end{bmatrix} \tag{3}$$

Reference space $S_r$ is a mathematical space, with genes in rows and axes as columns. $s_{ik}^{g_s}$ is the fold change of gene $i$ of the stress $k$. Stress $k$ itself is part of the stress group $g_s$. For instance, stress $k$ can be *P. infestans*, *Pseudomonas*, *Alternaria*, or *Golovinomyces orontii*, for all of which $g_s$ is "Biotic" stress. We can average over all columns, i.e. axes, corresponding to each stress group, to yield a mean reference space:

$$\overline{S_r} = \begin{bmatrix} \overline{s_{11}} & \overline{s_{12}} & \dots & \overline{s_{1q}} \\ \overline{s_{21}} & \overline{s_{22}} & \dots & \overline{s_{2q}} \\ \vdots & \vdots & \ddots & \vdots \\ \overline{s_{n1}} & \overline{s_{n2}} & \dots & \overline{s_{nq}} \end{bmatrix} \tag{4}$$

In equation 4, $\overline{s_{iq}} = \frac{1}{N} \sum_{z \in g_q} s_{iz}$.

We chose all *A. Thaliana* array samples that are measured by Affymetrix Arabidopsis ATH1 Genome Array as the main training data, for making the main reference space throughout the results section. The reason for this selection was twofold: firstly, with this selection, large enough samples, encompassing most stress groups, are included in the training data. And secondly, the reference space is exclusively calculated from Arabidopsis microarray data. Hence, demonstrating this reference is applicable on test data generated using RNA-seq or from another species, we can show how PhysioSpace can be used to translate among different species and platform technologies. Based on all *A. thaliana* array samples, we generated a reference space $S_r$, and a subsequent mean-space $\overline{S_r}$.
