## Supplemental File 2 for "Plant PhysioSpace: a robust tool to compare stress response across plant species"

### Supplement File 2: GO Analysis Results of the Reference Space

Esfahani et al.

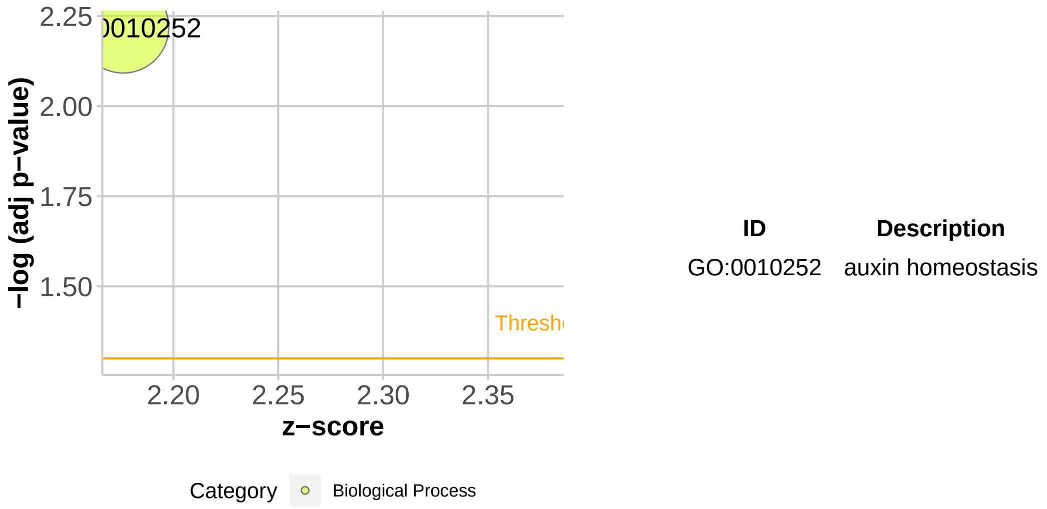


**Figure 1: GO Analysis of BioMone Stress from the Mean Stress Space.** In the plot, each enriched GO term is represented by a circle, with adjusted *p*-values as y-axis and enrichment ratio as x-axis. The size of the circle shows the size of the gene list of the corresponding GO term. And enrichment ratio here means the ratio between the actual number of differentially expressed genes and the expected, in each GO group. 5 most significant GO terms (or less in case less than 5 GO terms were significant.) are labeled on the plot and listed in a table on the right. The plot is generated using the GOplot package in R [1].

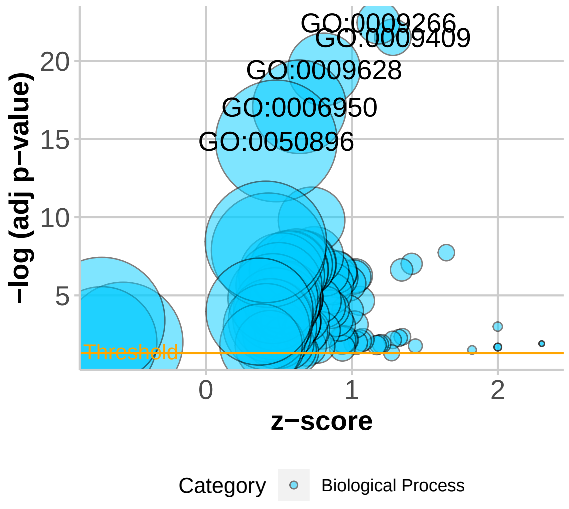


| ID | Description |
|---|---|
| GO:0009409 | response to cold |
| GO:0009266 | response to temperature stimulus |
| GO:0009628 | response to abiotic stimulus |
| GO:0006950 | response to stress |
| GO:0050896 | response to stimulus |

**Figure 2: GO Analysis of Cold Stress from the Mean Stress Space.** In the plot, each enriched GO term is represented by a circle, with adjusted $p$-values as y-axis and enrichment ratio as x-axis. The size of the circle shows the size of the gene list of the corresponding GO term. And enrichment ratio here means the ratio between the actual number of differentially expressed genes and the expected, in each GO group. 5 most significant GO terms (or less in case less than 5 GO terms were significant.) are labeled on the plot and listed in a table on the right. The plot is generated using the GOplot package in R [1].

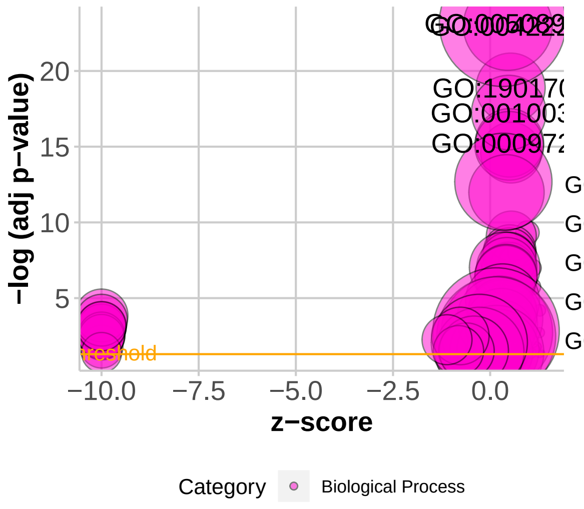


| ID | Description |
|---|---|
| GO:1901700 | response to oxygen−containing compoun |
| GO:0009725 | response to hormone |
| GO:0010033 | response to organic substance |
| GO:0042221 | response to chemical |
| GO:0050896 | response to stimulus |

**Figure 3: GO Analysis of Drought.Light Stress from the Mean Stress Space.** In the plot, each enriched GO term is represented by a circle, with adjusted $p$-values as y-axis and enrichment ratio as x-axis. The size of the circle shows the size of the gene list of the corresponding GO term. And enrichment ratio here means the ratio between the actual number of differentially expressed genes and the expected, in each GO group. 5 most significant GO terms (or less in case less than 5 GO terms were significant.) are labeled on the plot and listed in a table on the right. The plot is generated using the GOplot package in R [1].

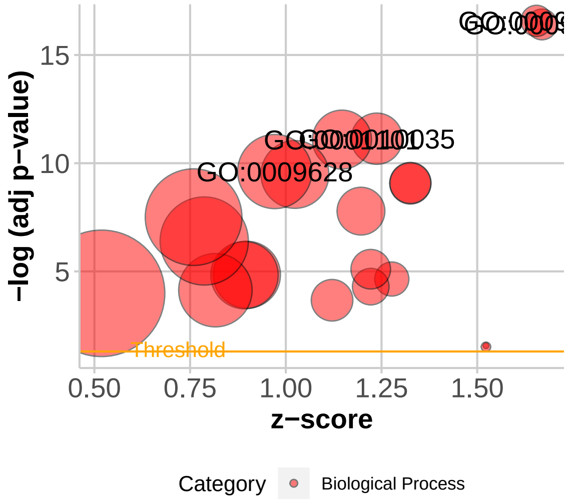


| ID | Description |
|---|---|
| GO:0009414 | response to water deprivation |
| GO:0009415 | response to water |
| GO:0010035 | response to inorganic substance |
| GO:0001101 | response to acid chemical |
| GO:0009628 | response to abiotic stimulus |

**Figure 4: GO Analysis of DrouSaTic Stress from the Mean Stress Space.** In the plot, each enriched GO term is represented by a circle, with adjusted $p$-values as y-axis and enrichment ratio as x-axis. The size of the circle shows the size of the gene list of the corresponding GO term. And enrichment ratio here means the ratio between the actual number of differentially expressed genes and the expected, in each GO group. 5 most significant GO terms (or less in case less than 5 GO terms were significant.) are labeled on the plot and listed in a table on the right. The plot is generated using the GOplot package in R [1].

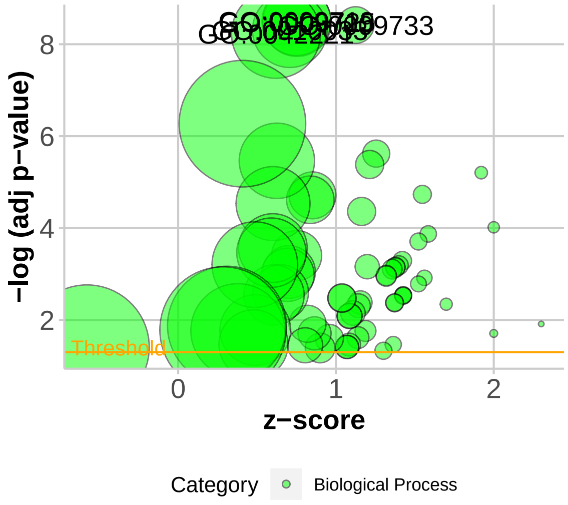


| ID | Description |
|---|---|
| GO:0009733 | response to auxin |
| GO:0009725 | response to hormone |
| GO:0009719 | response to endogenous stimulus |
| GO:0010033 | response to organic substance |
| GO:0042221 | response to chemical |

**Figure 5: GO Analysis of FarRed Stress from the Mean Stress Space.** In the plot, each enriched GO term is represented by a circle, with adjusted $p$-values as y-axis and enrichment ratio as x-axis. The size of the circle shows the size of the gene list of the corresponding GO term. And enrichment ratio here means the ratio between the actual number of differentially expressed genes and the expected, in each GO group. 5 most significant GO terms (or less in case less than 5 GO terms were significant.) are labeled on the plot and listed in a table on the right. The plot is generated using the GOplot package in R [1].

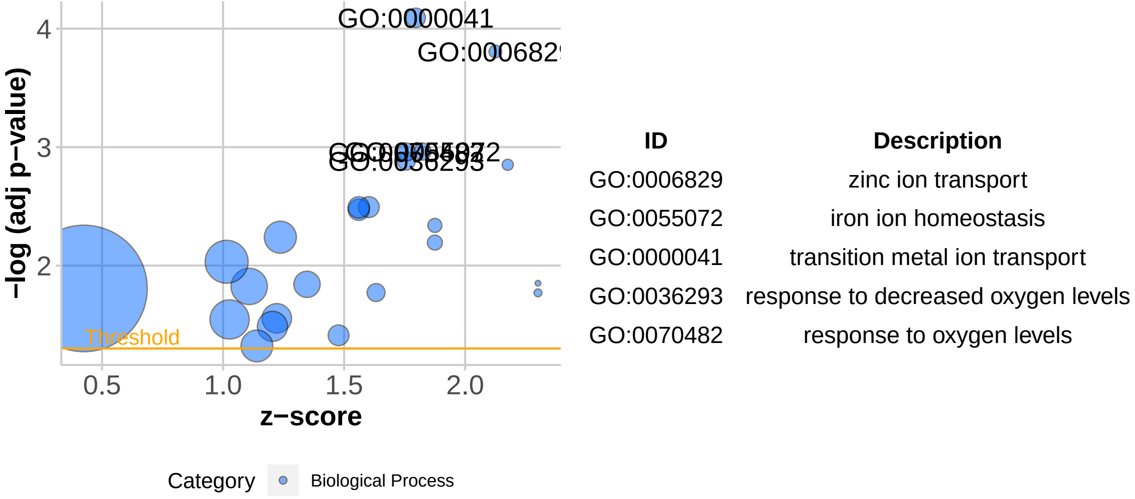


**Figure 6: GO Analysis of FeDeficiency Stress from the Mean Stress Space.** In the plot, each enriched GO term is represented by a circle, with adjusted $p$-values as y-axis and enrichment ratio as x-axis. The size of the circle shows the size of the gene list of the corresponding GO term. And enrichment ratio here means the ratio between the actual number of differentially expressed genes and the expected, in each GO group. 5 most significant GO terms (or less in case less than 5 GO terms were significant.) are labeled on the plot and listed in a table on the right. The plot is generated using the GOplot package in R [1].

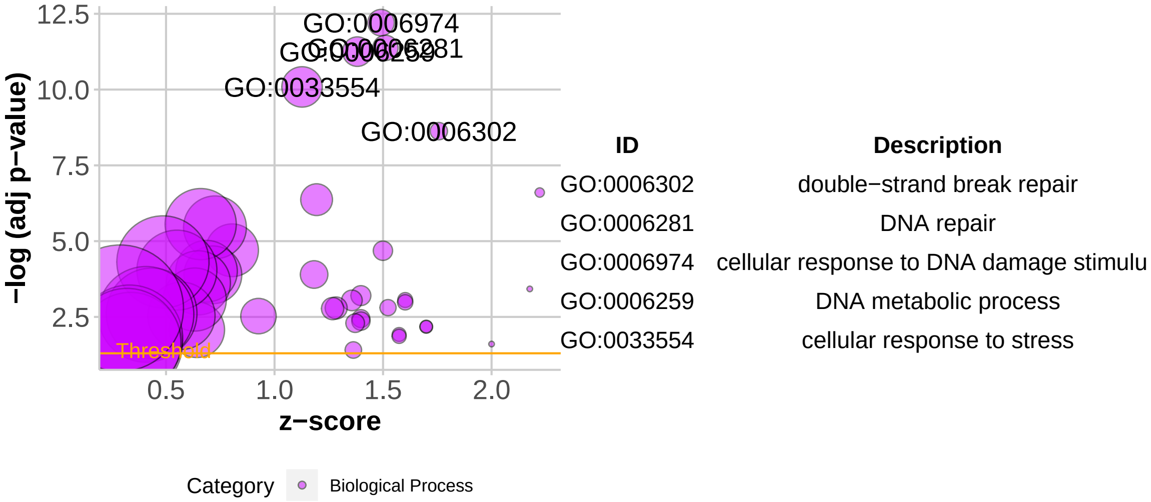


**Figure 7: GO Analysis of Genotoxic Stress from the Mean Stress Space.** In the plot, each enriched GO term is represented by a circle, with adjusted $p$-values as y-axis and enrichment ratio as x-axis. The size of the circle shows the size of the gene list of the corresponding GO term. And enrichment ratio here means the ratio between the actual number of differentially expressed genes and the expected, in each GO group. 5 most significant GO terms (or less in case less than 5 GO terms were significant.) are labeled on the plot and listed in a table on the right. The plot is generated using the GOplot package in R [1].

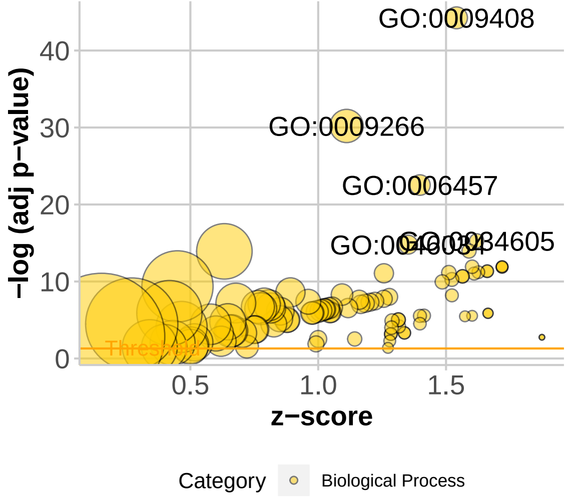


| ID | Description |
|---|---|
| GO:0034605 | cellular response to heat |
| GO:0009408 | response to heat |
| GO:0006457 | protein folding |
| GO:0046034 | ATP metabolic process |
| GO:0009266 | response to temperature stimulus |

**Figure 8: GO Analysis of Heat Stress from the Mean Stress Space.** In the plot, each enriched GO term is represented by a circle, with adjusted $p$-values as y-axis and enrichment ratio as x-axis. The size of the circle shows the size of the gene list of the corresponding GO term. And enrichment ratio here means the ratio between the actual number of differentially expressed genes and the expected, in each GO group. 5 most significant GO terms (or less in case less than 5 GO terms were significant.) are labeled on the plot and listed in a table on the right. The plot is generated using the GOplot package in R [1].

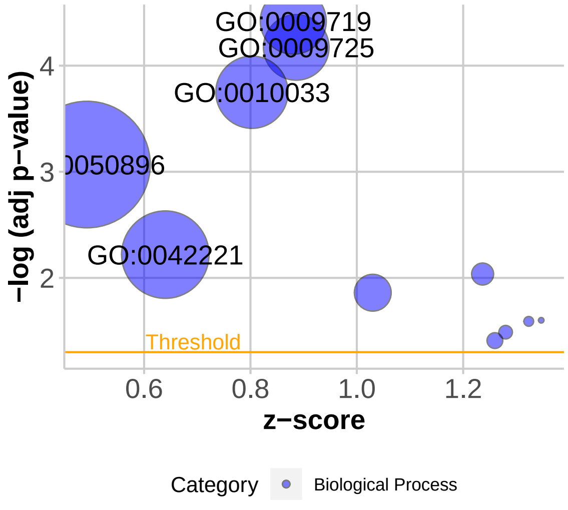


| ID | Description |
|---|---|
| GO:0009725 | response to hormone |
| GO:0009719 | response to endogenous stimulus |
| GO:0010033 | response to organic substance |
| GO:0042221 | response to chemical |
| GO:0050896 | response to stimulus |

**Figure 9: GO Analysis of Herbicide Stress from the Mean Stress Space.** In the plot, each enriched GO term is represented by a circle, with adjusted $p$-values as y-axis and enrichment ratio as x-axis. The size of the circle shows the size of the gene list of the corresponding GO term. And enrichment ratio here means the ratio between the actual number of differentially expressed genes and the expected, in each GO group. 5 most significant GO terms (or less in case less than 5 GO terms were significant.) are labeled on the plot and listed in a table on the right. The plot is generated using the GOplot package in R [1].

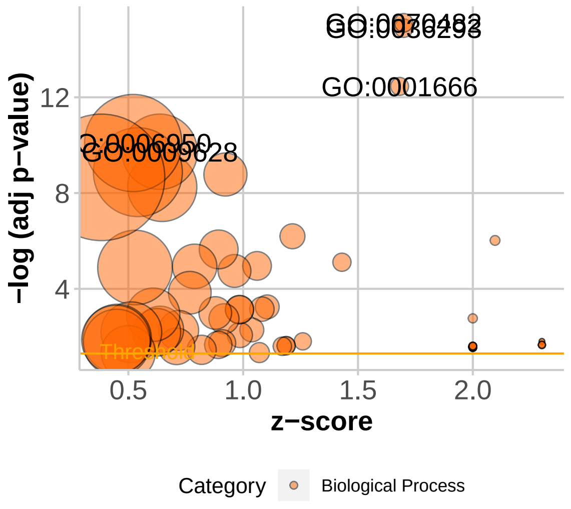


| ID | Description |
|---|---|
| GO:0036293 | response to decreased oxygen levels |
| GO:0070482 | response to oxygen levels |
| GO:0001666 | response to hypoxia |
| GO:0009628 | response to abiotic stimulus |
| GO:0006950 | response to stress |

**Figure 10: GO Analysis of Hypoxia Stress from the Mean Stress Space.** In the plot, each enriched GO term is represented by a circle, with adjusted $p$-values as y-axis and enrichment ratio as x-axis. The size of the circle shows the size of the gene list of the corresponding GO term. And enrichment ratio here means the ratio between the actual number of differentially expressed genes and the expected, in each GO group. 5 most significant GO terms (or less in case less than 5 GO terms were significant.) are labeled on the plot and listed in a table on the right. The plot is generated using the GOplot package in R [1].

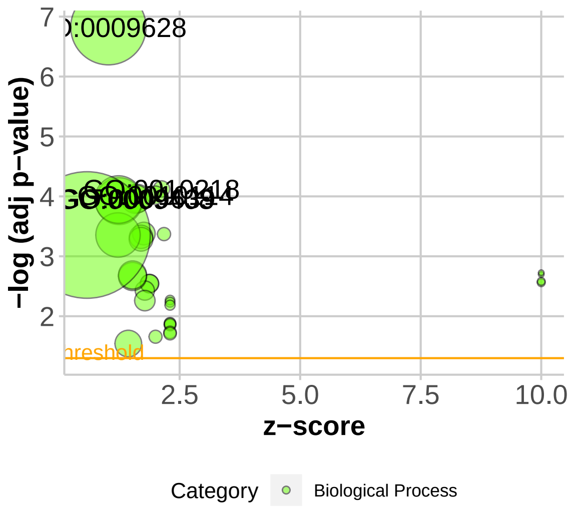


| ID | Description |
|---|---|
| GO:0010218 | response to far red light |
| GO:0010114 | response to red light |
| GO:0009639 | response to red or far red light |
| GO:0009408 | response to heat |
| GO:0009628 | response to abiotic stimulus |

**Figure 11: GO Analysis of LighUV Stress from the Mean Stress Space.** In the plot, each enriched GO term is represented by a circle, with adjusted $p$-values as y-axis and enrichment ratio as x-axis. The size of the circle shows the size of the gene list of the corresponding GO term. And enrichment ratio here means the ratio between the actual number of differentially expressed genes and the expected, in each GO group. 5 most significant GO terms (or less in case less than 5 GO terms were significant.) are labeled on the plot and listed in a table on the right. The plot is generated using the GOplot package in R [1].

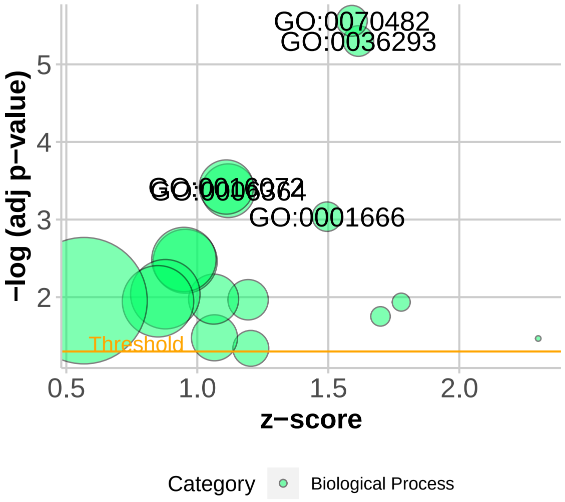


| ID | Description |
|---|---|
| GO:0036293 | response to decreased oxygen levels |
| GO:0070482 | response to oxygen levels |
| GO:0001666 | response to hypoxia |
| GO:0006364 | rRNA processing |
| GO:0016072 | rRNA metabolic process |

**Figure 12: GO Analysis of LowPH Stress from the Mean Stress Space.** In the plot, each enriched GO term is represented by a circle, with adjusted $p$-values as y-axis and enrichment ratio as x-axis. The size of the circle shows the size of the gene list of the corresponding GO term. And enrichment ratio here means the ratio between the actual number of differentially expressed genes and the expected, in each GO group. 5 most significant GO terms (or less in case less than 5 GO terms were significant.) are labeled on the plot and listed in a table on the right. The plot is generated using the GOplot package in R [1].

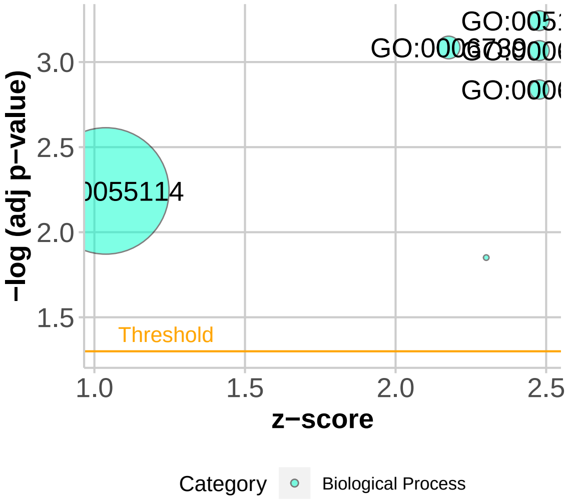


| ID | Description |
|---|---|
| GO:0006098 | pentose−phosphate shunt |
| GO:0006740 | NADPH regeneration |
| GO:0051156 | glucose 6−phosphate metabolic process |
| GO:0006739 | NADP metabolic process |
| GO:0055114 | oxidation−reduction process |

**Figure 13: GO Analysis of Nitrogen Stress from the Mean Stress Space.** In the plot, each enriched GO term is represented by a circle, with adjusted $p$-values as y-axis and enrichment ratio as x-axis. The size of the circle shows the size of the gene list of the corresponding GO term. And enrichment ratio here means the ratio between the actual number of differentially expressed genes and the expected, in each GO group. 5 most significant GO terms (or less in case less than 5 GO terms were significant.) are labeled on the plot and listed in a table on the right. The plot is generated using the GOplot package in R [1].

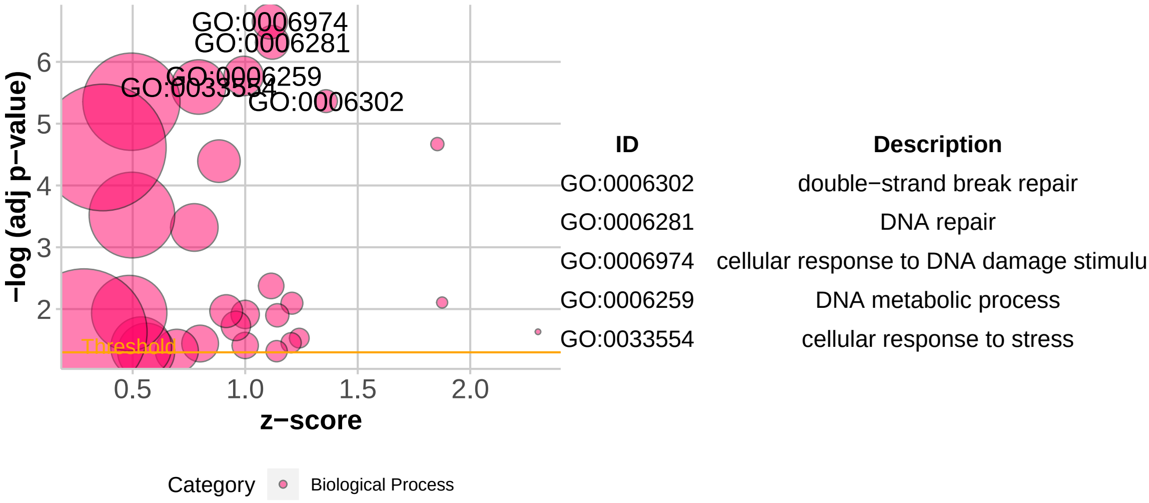


**Figure 14: GO Analysis of Radiation Stress from the Mean Stress Space.** In the plot, each enriched GO term is represented by a circle, with adjusted $p$-values as y-axis and enrichment ratio as x-axis. The size of the circle shows the size of the gene list of the corresponding GO term. And enrichment ratio here means the ratio between the actual number of differentially expressed genes and the expected, in each GO group. 5 most significant GO terms (or less in case less than 5 GO terms were significant.) are labeled on the plot and listed in a table on the right. The plot is generated using the GOplot package in R [1].

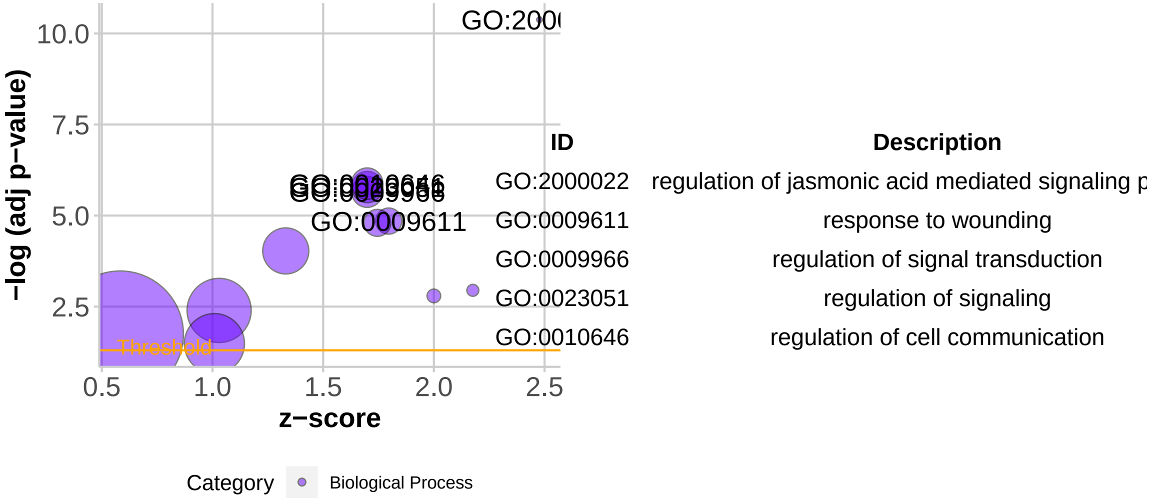


**Figure 15: GO Analysis of Wounding Stress from the Mean Stress Space.** In the plot, each enriched GO term is represented by a circle, with adjusted $p$-values as y-axis and enrichment ratio as x-axis. The size of the circle shows the size of the gene list of the corresponding GO term. And enrichment ratio here means the ratio between the actual number of differentially expressed genes and the expected, in each GO group. 5 most significant GO terms (or less in case less than 5 GO terms were significant.) are labeled on the plot and listed in a table on the right. The plot is generated using the GOplot package in R [1].
