## Supplementary figures and images for "Plant PhysioSpace: a robust tool to compare stress response across plant species"

### Supplemental Figure S1

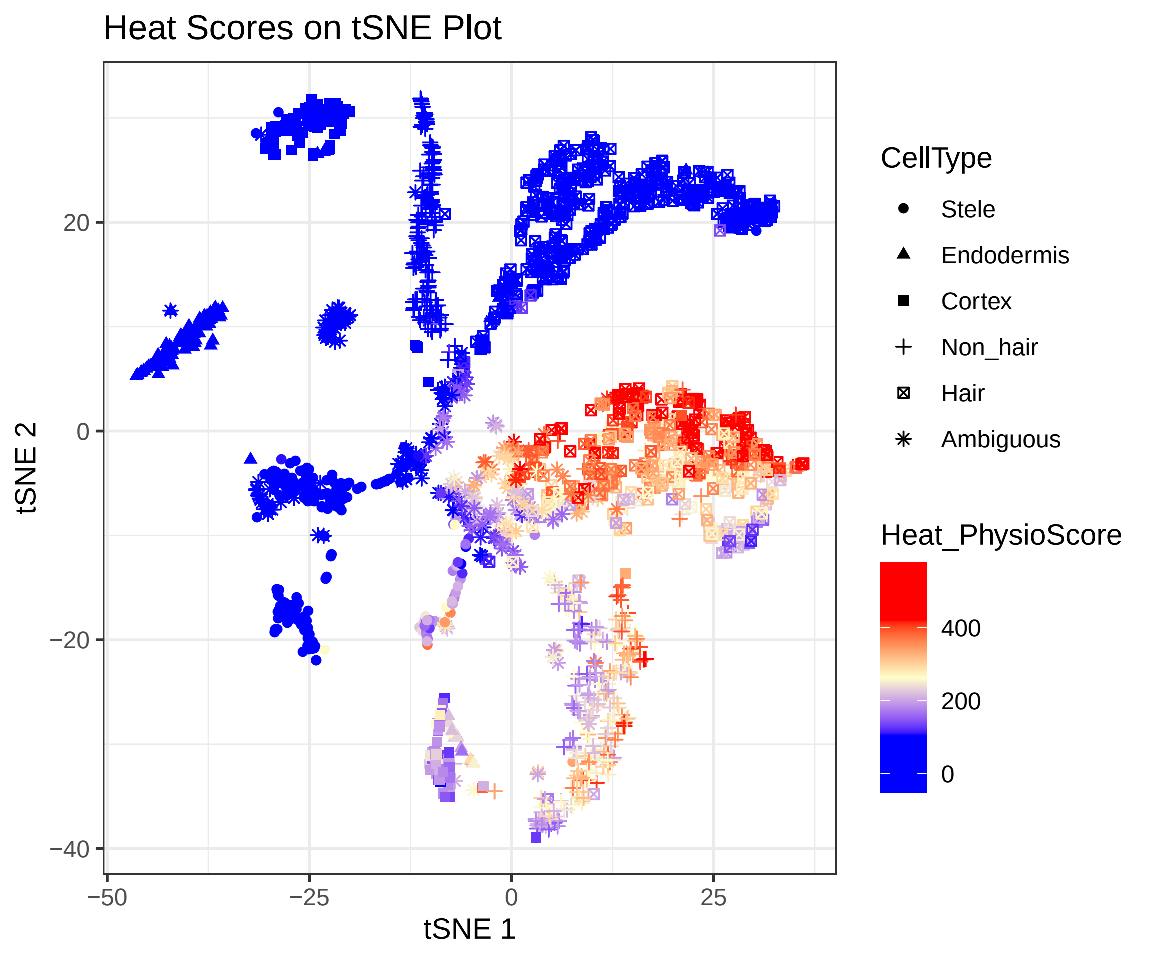

### Supplemental Figure S2

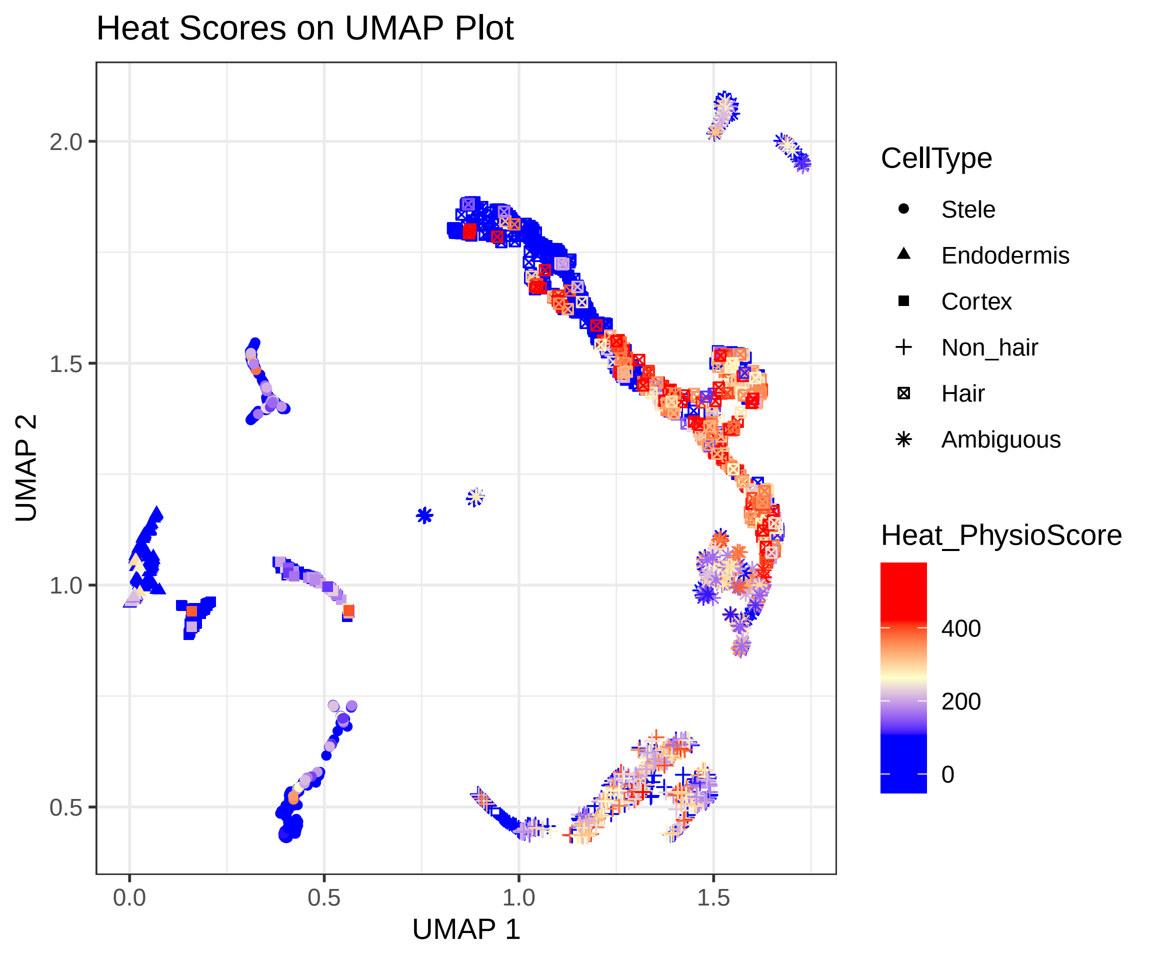

### Supplemental Figure S3

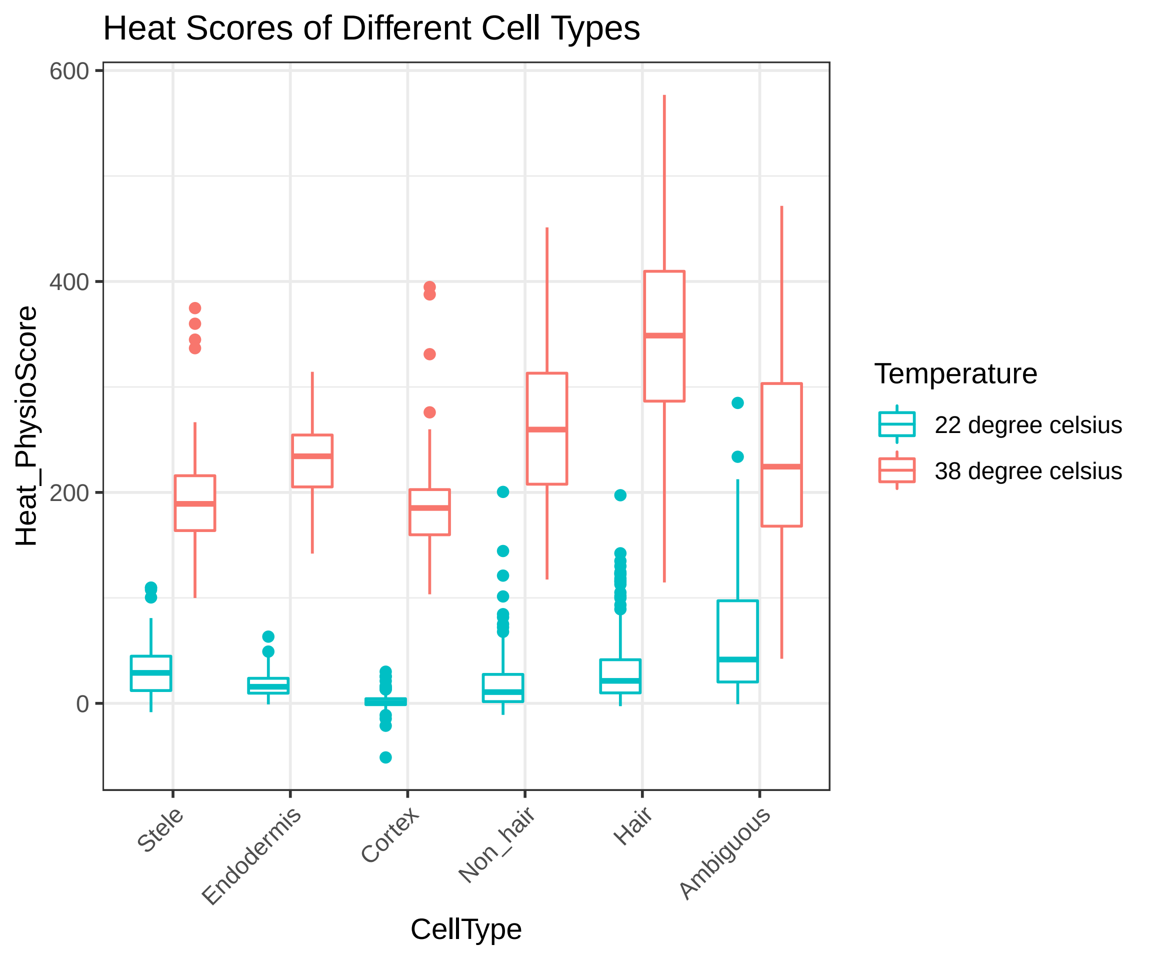
